# Disorder or a new order: how climate change affects phenological variability

**DOI:** 10.1101/2021.10.08.463688

**Authors:** Michael Stemkovski, James R. Bell, Elizabeth R. Ellwood, Brian D. Inouye, Hiromi Kobori, Sang Don Lee, Trevor Lloyd-Evans, Richard B. Primack, Barbara Templ, William D. Pearse

**Affiliations:** Department of Biology & Ecology Center, Utah State University, Logan, UT, USA; Rocky Mountain Biological Laboratory, Crested Butte, CO, USA; Rothamsted Research, West Common, Harpenden, Hertfordshire, UK; Natural History Museum of Los Angeles County, Los Angeles, California, USA; iDigBio, Florida Museum of Natural History, University of Florida, Gainesville, Florida, USA; Department of Biological Science, Florida State University, Tallahassee, FL, USA; Tokyo City University, Tamazutsumi, Setagaya, Tokyo, Japan; Department of Environmental Science and Engineering, Ewha Womans University, Seoul, Republic of Korea; Manomet Observatory, Manomet, MA, USA; Department of Biology, Boston University, Boston, MA, USA; data-analysis OG, Vienna, Austria; Department of Life Sciences, Imperial College London, Silwood Park Campus, Ascot, Berkshire, UK

**Author notes:** Author emails: M.S., J.R.B., E.R.E, B.D.I., H.K., S.D.L, T.L.E., R.B.P., B.T., W.D.P.

## Abstract

Advancing spring phenology is a well-documented consequence of anthropogenic climate change, but it is not well understood how climate change will affect the variability of phenology year-to-year. Species’ phenological timings reflect adaptation to a broad suite of abiotic needs (e.g. thermal energy) and biotic interactions (e.g. predation and pollination), and changes in patterns of variability may disrupt those adaptations and interactions. Here, we present a geographically and taxonomically broad analysis of phenological shifts, temperature sensitivity, and changes in inter-annual variance encompassing nearly 10,000 long-term phenology time-series representing over 1,000 species across much of the northern hemisphere. We show that early-season species in colder and less seasonal regions were the most sensitive to temperature change and had the least variable phenologies. The timings of leaf-out, flowering, insect first-occurrence, and bird arrival have all shifted earlier and tend to be less variable in warmer years. This has led leaf-out and flower phenology to become moderately but significantly less variable over time. These simultaneous changes in phenological averages and the variation around them have the potential to influence mismatches among interacting species that are difficult to anticipate if shifts in average are studied in isolation.

## Introduction

Shifts in the timing of life-cycle events (phenology) have occurred as a result of changes in climate, and while there has been a general trend of species in seasonal regions advancing their spring phenology over the last few decades due to anthropogenic climate warming (Parmesan et al., 2003), species vary in their phenological responses to inter-annual climatic variation. Species’ phenological sensitivity varies based on trophic level (Thackeray et al., 2016), and insects are thought to be able to track climatic cues more closely than other groups of animals and plant (Cohen et al., 2018). Species’ traits can influence phenological sensitivity within many taxonomic groups including plants (Konig et al., 2018), insects (Diamond et al., 2011), and birds (Butler, 2003). Phenological shifts have been more pronounced in early season species (CaraDonna et al., 2014; Mulder et al., 2017) occupying colder regions (Roslin et al., 2021) and higher latitudes (Parmesan, 2007), likely due to the faster pace of climate change in the upper northern hemisphere (Burrows et al., 2011) and stronger selection for plasticity (Lindestad et al., 2019) in those areas. Some species have also shown decreases in phenological sensitivity to temperature variation in warmer years as they reach the limits of their historical climate conditions, producing non-linear temperature-phenology relationships (Iler et al., 2013; Mulder et al., 2017). While these complexities alone make it difficult to predict how species’ phenologies will change in the future, it is also unclear whether climate change is making phenology inherently more or less variable and predictable. Such changes in variability are not just of academic concern, particularly if they affect the reliability of species’ interactions that drive key ecosystem services such as pollination for agriculture.

The majority of phenological research has focused on changes in the mean of events such as onset and peak over time (phenological shifts) or in response to yearly climatic variation (phenological sensitivity). Some studies have also shown changes in within-season (*intra*-annual) variance due to climate change (Zohner et al., 2018), but few have investigated whether, or in what ways, the variance of phenological events across years (*inter*-annual) is being affected by climate change. Most studies have assumed constant inter-annual variance, and some have checked and accounted for heteroscedasticity in time-series residuals (e.g., Bartomeus et al., 2011; Wadgymar et al., 2018) but have not made it a focus of study. There is reason to think that inter-annual variance in phenology might be changing, as there have been recent, geographically heterogeneous changes in inter-annual temperature variance (Liu et al., 2020). Further, decreased sensitive to temperature variation (Mulder et al., 2017), chilling requirements in plants (Fu et al., 2015), and physiological development time requirements between phenophases (Ettinger et al., 2018; Primack, 1987) may produce patterns of phenological variance that are different from the variance of their cues. To our knowledge, the studies that have examined inter-annual phenology variance on a broad scale, using citizen science data (Pearse et al., 2017) and satellite imagery (Liu et al., 2020), have found increases in variance over time, though such changes might also be influenced by changes in monitoring schemes or community composition over time (de Keyzer et al., 2017).

The scarcity of research attention does not reflect a lack of importance, as changes in phenological variance can have consequences for the temporal synchrony of interacting species. An increase in phenological variability may hamper the ability of dependent species to track the moving target of other species’ changing phenology if the species track different climatic cues or have different sensitivities to the same cues. Variation in phenological overlap affects the strength of interactions between co-occurring species (Tiusanen et al., 2020), so there might be immediate consequences for species’ fitness and coexistence. Extreme inter-annual phenological variation in the overlap of interacting species may even lead to local extirpation (Patterson et al., 2020). While phenological mismatches resulting in short-term fitness losses may be followed by evolutionary adaptation in plasticity that corrects the mismatch (Visser et al., 2019), this adaptation may be less likely to occur if the phenological fitness landscape becomes less predictable (Leung et al., 2020). If environments become extremely unpredictable, species may even adapt bet-hedging strategies rather than maintain plasticity (Botero et al., 2015). Beyond predictability, changes in variance can even influence mean shifts in phenology by interacting with lagged effects of temperature on leaf and flower primordia in previous years (Mulder et al., 2017). From a cultural standpoint, changes in the phenological mean and variance affect how possible it is to plan ahead for events such as flowering festivals and autumn leaf-viewing seasons (Allen et al., 2014).

In the present study, we examine nearly 10,000 time-series datasets of plant, insect, and bird phenology to determine the general patterns of how climate change is affecting both phenological means and variance. To do this, we specify four metrics of change (Figure 1): mean change over many years (mean shift), inter-annual mean changes due to climate variability (mean sensitivity), variance change over years (variance shift), and inter-annual variance changes due to climate variability (variance sensitivity). We identify the regional climatic drivers of shifts and sensitivity, the effect of phenological position (how early in the season a phenophase occurs), and differences among taxonomic groups. Finally, we examine the influence of functional traits on shifts and sensitivity within groups. We confirm results from previous studies, that early-season species are on average more sensitive to temperature variation and that phenology in regions with colder climates is advancing at the greatest rate. Contrary to previous studies, we do not find evidence of increasingly variable inter-annual spring phenology in any taxonomic group, and in fact detect decreases in variability in warmer years in all groups and a slight overall decrease in variability over time in leaf-out and flowering. We argue that these patterns indicate that recent climate change in the northern hemisphere is shifting spring earlier but is not necessarily making it more erratic. This suggests that we might be more able to predict and prepare for the consequences of ecological rearrangements resulting from climate change.

**Figure 1.**
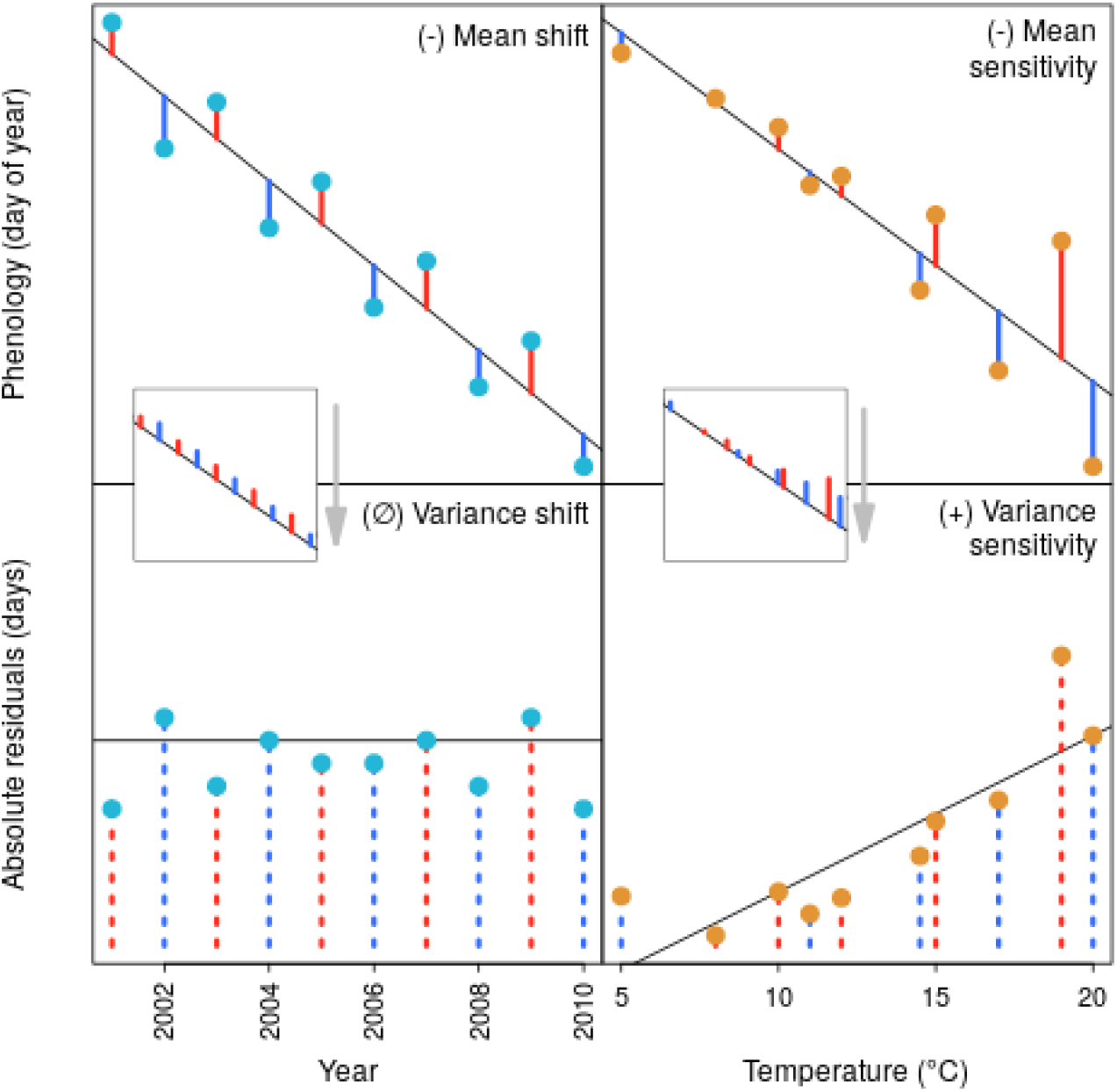
Conceptual demonstration of the four phenology shift and sensitivity metrics. Phenological mean shift and mean sensitivity (top panels) are defined as the slope of the relationship between the day of year on which a phenophase was observed and the year or temperature, respectively, associated with that observation. Shift and sensitivity of variance (bottom panels) are then computed as the slope of the absolute residuals versus the year/temperature. Teal points represent yearly data, and orange ones represent data relating to inter-annual temperature variation. Red lines indicate positive residuals, blue lines represent negative residuals, and dashed lines represent absolute residuals. The middle, pop-out subfigures highlight the intermediate process of taking the absolute value of the residuals from the mean regression in the top panels in order to compute variance changes in the bottom panels. This hypothetical example demonstrates a case in which mean phenology is shifting earlier (top-left), is earlier in warmer years (top-right), variance is not shifting over time (bottom-left), but variance is greater in warmer years (bottom-right).

## Methods

In order to determine how phenological means and variance are shifting over time and how sensitive they are to inter-annual climate variation, we pooled data from eight long-term monitoring schemes, calculated four phenology metrics (Figure 1) for individual time-series within these datasets, and modeled the resulting trends using regional climate and species characteristics. All analyses were done in R version 3.6.3 (R Core Team, 2020). Data management was done using the R-package *data*.*table* (Dowle et al., 2021), quantile regression was done using *quantreg* (Koenker, 2021), and data visualization was aided by *visreg* (Breheny et al., 2015), *rnatualearth* (South, 2017), *sf* (Pebesma, 2021), and *cowplot* (Wilke, 2020). Scripts to reproduce analysis are available online (https://github.com/stemkov/pheno_variance) and in the supplementary materials.

### Phenology data

We performed a broad search of long-term phenological datasets across terrestrial taxa. We included datasets with time-series spanning longer than 10 years, extending at least past the year 2000, and for which measurements were made repeatedly by experts at fixed locations. We included eight sources: Korean meteorological stations (Ibáñez et al., 2010; Kim et al., 2021), Japanese meteorological stations (Doi et al., 2008; Ibáñez et al., 2010), the NECTAR network (Cook et al., 2012), the Rocky Mountain Biological Lab (RMBL; CaraDonna et al., 2014; Inouye, 2008), the Manomet Observatory bird monitoring station (Lloyd-Evans et al., 2004; Stegman et al., 2017), the Rothamsted aphid trap network (Bell et al., 2015), the Chronicles of Nature Calendar (Ovaskainen et al., 2020), and the Pan-European Phenology network (Templ et al., 2018). Further details on these data sources are provided in Table S7.1.

We focused our analysis on plant leaf-out, the onset of plant flowering, the first appearance of adult insects, and the first arrival of migratory birds. We refer to these four groups henceforth as phenophase groups. We used first-observation dates as the measure of phenological onset for most datasets due to the unavailability of continuous abundance records and in most of the datasets. The Manomet and RMBL datasets include seasonal abundance time-series, so we were able to more precisely estimate phenological onset using a Weibull estimator (Pearse et al., 2017). We note that first-occurrence dates may not reflect shifts in the peak or duration of phenophases (Inouye et al., 2019), but we did not investigate these due to limitations of the present datasets. We performed systematic quality assurance and excluded time-series based on the following five conditions that were likely to lead to erroneous shift and sensitivity calculations. Of the 15,930 total time-series that we evaluated, we (1) excluded 10 that contained gaps in observations that made up more than three quarters of the time-series range. (2) We excluded 30 time-series that were unusable due to ambiguous data recording schemes in which some phenophases were recorded in January and some in December, but the year was unclear. (3) 89 time-series were excluded due to potentially unreliable estimates flagged by the Weibull method implementation, with estimates not matching up to their confidence interval range (see Smith, 1987). (4) Unresolvable data entry errors were identified in 170 time-series when there were extreme outliers or discontinuous data clusters that might have been caused by swapping days and months in data entry. These cases were flagged using model-based clustering (Fraley et al., 2012) with a conservative model selection threshold of *BIC=25*, and potential cases of clustering were checked visually. Lastly (5), 618 time-series with a total range of observations greater than three months were excluded due to likely aseasonal dynamics. Much of the NECTAR data was removed due to fewer than 10 years of recent observations at most sites. To avoid problems with pseudoreplication due to co-located or spatially clustered sites in the CNC and PEP datasets, we picked the co-located CNC sites with the most records, and in the PEP data selected the sites with the most records for each decimal coordinate rounded to the nearest whole (which is c. 55km apart in Europe). This selection process left 288 of the 354 sites in the CNC data and 360 of the 15,183 locations in the PEP data.

### Climate data

We obtained geographically precise historical climate data from the TerraClimate product (Abatzoglou et al., 2018), which provides monthly maximum temperature estimates at a ∼4km resolution globally from 1958 to 2018. To match this data product, we did not consider data earlier than 1958 or after 2018. To calculate a series of relevant yearly temperatures for each species/phenophase/site time-series, we identified the median month in which the phenophase occurred across the entire time-series, and extracted the average maximum temperature in that month across all years of the time-series. Because these temperature measurements are used to estimate mean and variance sensitivity (detailed below), we note that extracting the temperature at the median months may produce conservative estimates compared to approaches characterizing entire climate sensitivity profiles (Thackeray et al., 2016), though by using a fixed integration period length of one month, we ensure comparability across datasets (following Keenan et al., 2020). We also note that temperature sensitivity is often calculated using degree-day models, though a comparison of these models against a simple linear regression approach similar to what we implemented showed that they provide similar results (Basler, 2016). We summarized the regional climate of sites with two metrics: seasonality and mean temperature. We defined seasonality as the mean annual temperature range (following Cook et al., 2012) in every year between 1958 and 2018, and mean temperature simply as the mean of monthly temperatures across all months in all years.

### Trait data

We obtained data on plant traits from the BIEN database (Maitner et al., 2018). We limited our selection of plant traits to those for that we had over 50% coverage and those which we hypothesized could be influential to leaf or flower phenology (Díaz et al., 2004): whole plant growth form, height, specific leaf area (SLA), and seed mass. We grouped whole plant growth form into five categories: trees, shrubs, herbs, grasses, and dependents. Herbs contained plants classified as forbs, ferns, hemicryptophytes, and geophytes, while dependents contained vines, epiphytes, hemiepiphytes, lianas, parasites, and other climbing plants. We excluded aquatic plants and cacti. We obtained data on bird body-mass and diet from the EltonTraits database (Wilman et al., 2014). In order to maximize the generality of the bird trait analysis and to create groups with comparable representation, we grouped herbivores, granivores, and frugivores into one “herbivore” group and combined those feeding primarily on invertebrates, vertebrates, and scavengers into one “carnivore” group. This resulted in three broad diet groups of herbivores, omnivores, and carnivores. We note that, in addition to these traits, migration distance may explain trends in bird phenology (Butler, 2003; Miller-Rushing et al., 2008) but this is not included in the present study due to limited data availability.

### Calculation of shifts and sensitivities

We calculated the rates of phenological mean shift for each species/site time-series by modeling the day-of-year (DOY) on which a phenophase was recorded as a linear function of year. Mean sensitivity was similarly calculated with DOY as a linear function of the monthly temperature associated with that observation. We calculated variance shifts and sensitivities by estimating the variance function using absolute residuals (following Davidian & Carroll, 1987). To estimate change in the standard deviation of the error function, we computed the absolute value of the residuals (|*R*_*i*_|) from the mean shift and sensitivity models and modeled the absolute residuals as a function of year and temperature, respectively, using quantile regression (Koenker et al., 2001) with *τ* ≈ *0*.*6827* (corresponding to the proportion of the absolute residual distribution found within one standard deviation of zero). Calculation of standard errors and significance testing for the quantile regressions were done using bootstrapping with the default xy-pairs method and 200 replicates. The calculation of the four metrics is visualized in Figure 1, and we performed a simulation study to confirm that the absolute residual approach is unbiased at detecting variance shifts (Supplement 1). We also tested for the effects of potential non-linearity on mean and variance change calculations (Supplement 2).

### Analysis of trends

In order to determine the drivers of phenological mean shifts, temperature sensitivity, and variance changes, we performed several analyses on the estimated rates of shifts. First, we investigated whether regional climate (long-term seasonality and mean annual temperature) and phenological position (how early in the season a species’ phenophase typically occurs relative to others at the same site) predicted the magnitude of shifts, and whether different phenophase groups (leaves, flowers, insects, and birds) have all shifted similarly. To do this, we constructed four linear mixed effects models (Bates et al., 2020) with seasonality, mean temperature, phenological position, and phenophase group as additive fixed effects, and species and sites within datasets as categorical random effects. To propagate uncertainty of shift estimates due to variable time-series lengths and correlation strength, we weighted the regressions by the inverse of the standard errors of the *μ* and *σ* coefficients. In order to compare the effects of continuous and categorical predictors and to assess the relative importance of coefficient estimates, we centered and scaled the predictor variables by 0.5 standard deviations (Gelman, 2008), and tested for fixed-effect term statistical significance (i.e., coefficients different from 0) using the *lmerTest* R-package (Kuznetsova et al., 2017).

To determine whether traits played a role in mean or variance shifts, we performed three secondary analyses on subsets of the flower, leaf, and bird data, each with the same random effects structure as in the model above. First, we tested whether four plant traits predicted shifts in flower phenology, with whole plant growth form, height, seed mass, and SLA as additive fixed effects. We conducted this analysis only for flowering phenology data because we obtained sparse data on leaf phenology for every growth form except shrubs and trees. In this and all subsequent models, we estimated a a reference intercept (dependents in the plant traits model, shrubs in the plant phenophase model, and carnivores in the bird traits model) and compared groups as contrasts from that intercept because we were interested in whether shifts varied significantly among groups. We then investigated whether flower and leaf phenology exhibited different shifts and whether there was an interaction with growth form for a subset of the data from shrubs, trees, and herbs, with growth form and phenophase as interacting fixed effects. Lastly, we analyzed the effect of body-mass and diet type (herbivore, omnivore, and carnivore) on phenological trends in birds by modeling the four metrics as functions of diet type interacting with the log_e_ body-mass of each bird species.

## Results

We analyzed 9,725 time-series with a median length of 36 years, representing 350,889 total phenological onset observations. The data were comprised of 2,399 leaf-out, 5,377 first flowering, 985 bird arrival, and 964 insect first-occurrence time-series. These data represented 1,048 species across 423 unique sites, with 801 plant, 168 bird, and 79 insect species (Figure 2b). The study sites were widely distributed across 24 countries in the temperate regions of the Northern Hemisphere and encompassed a wide climatic range, with mean annual temperature ranging from -1.9°C to 30.8°C, and the strength of seasonality ranging from a 3.5°C to 54.1°C difference between summer and winter temperature (Figure 2a).

**Figure 2.**
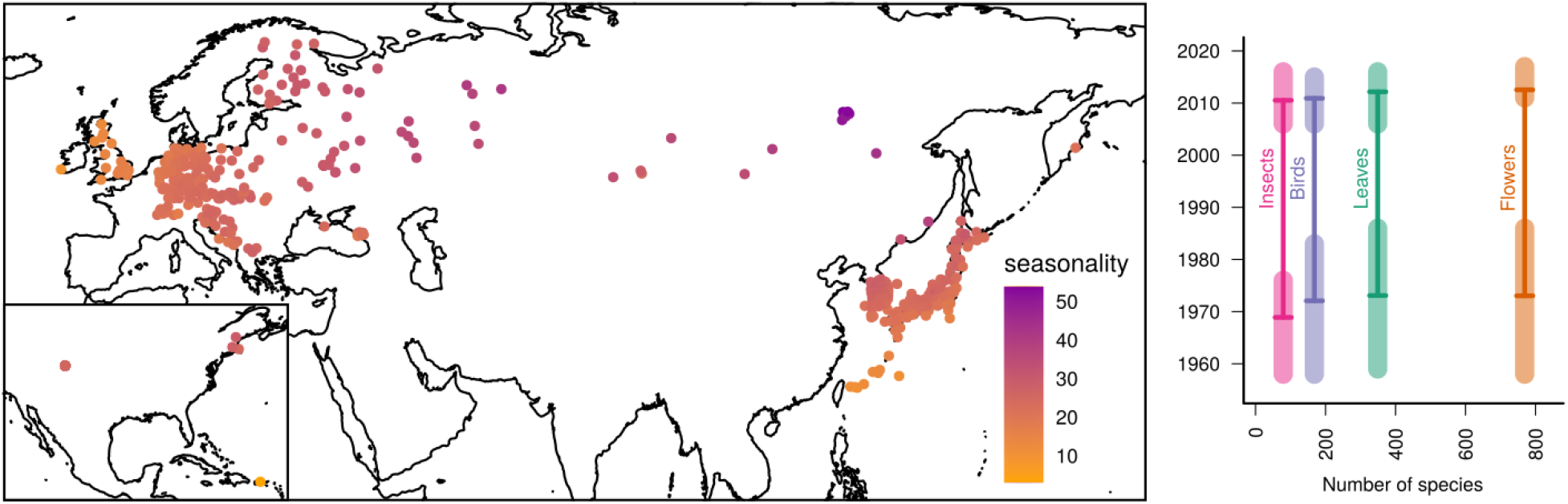
Spatial distribution of phenological data sources. Long-term phenological observation data has mostly been conducted in the temperature and boreal parts of the northern hemisphere, but the data used in this study are widely distributed and span a large gradient of regional climates (left panel). Yellow points represent sites with the least pronounced seasonal temperature differences, while purple ones represent the most seasonal sites. Seasonality is calculated as the annual mean temperature (°C) difference between the warmest and coldest months at each location. Most of the available phenological data is on plant phenophases, but the duration of time-series in the present dataset is roughly equal across taxonomic groups (right panel). Vertical bold lines represent the median duration of time-series for each phenophase group, with horizontal dashes representing the median start and end dates. The shaded bars around the horizontal dashes represent the first and third quartiles of the start and end years of the time-series.

We observed substantial variability in the strength and direction of mean shifts, mean sensitivity, variance shifts, and variance sensitivity. Across all phenophase groups, phenology advanced by 1.63 day/decade (mean shift; t_9722_ = -39.88, p < 0.001) and phenology was earlier in warmer years by 1.82 days/°C (mean sensitivity; t_9722_ = -96.5, p < 0.001; Figure 3a). Phenology became less variable in warmer years at a rate of 0.21 days/°C (variance sensitivity; t_9722_ = -17.24, p < 0.001; Figure 3b) and became less variable by 0.24 days/decade overall (variance shift; t_9722_ = -9.51, p < 0.001). For example, the variance of bigleaf hydrangea (*Hydrangea macrophylla*) flowering onset decreased by 0.81 days/°C on average across 86 sites, that of Norway maple (*Acer plantanoides*) decreased by 0.6 days/°C across 59 sites, and that of European blueberry (*Vaccinium myrtillus*) decreased by 1.35 days/decade across 23 sites. The degree of temperature sensitivity significantly predicted the shift in mean phenology over time (t_9721_ = 35.344, p < 0.001, R^2^ = 0.11; Figure 4), with greater sensitivity to temperature resulting in greater shifts toward earlier phenology over time.

**Figure 3.**
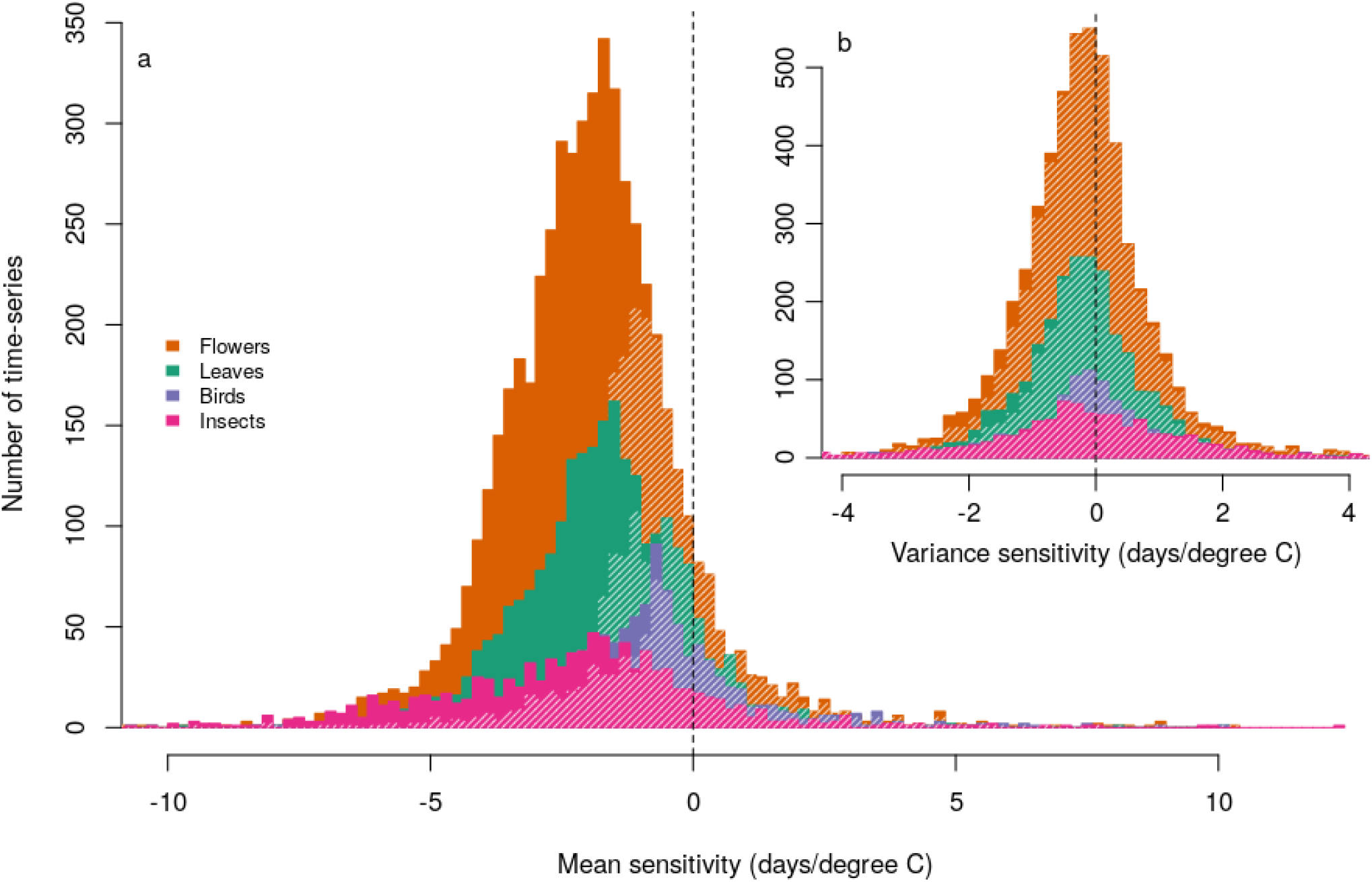
Phenological onset dates tend to be earlier and less variable in warmer years. The majority of all phenological groups (90% of flowers, 91% of leaves, 75% of birds, and 87% of insects) advanced their mean spring phenology in response to increased temperature (panel a). Phenology became less variable in warmer years, with 60% of time-series overall decreasing in variance (panel b). Time-series with individual slope estimates not significantly different from zero are shaded with white, and some of the data are obscured due to overlapping histograms. The plotting range is narrowed slightly to show the distributions more clearly, so 4 (<0.1%) points are excluded on the left of panel a, 68 points (0.6%) on the left of panel b, and 34 points (0.3%) on the right of panel b.

**Figure 4.**
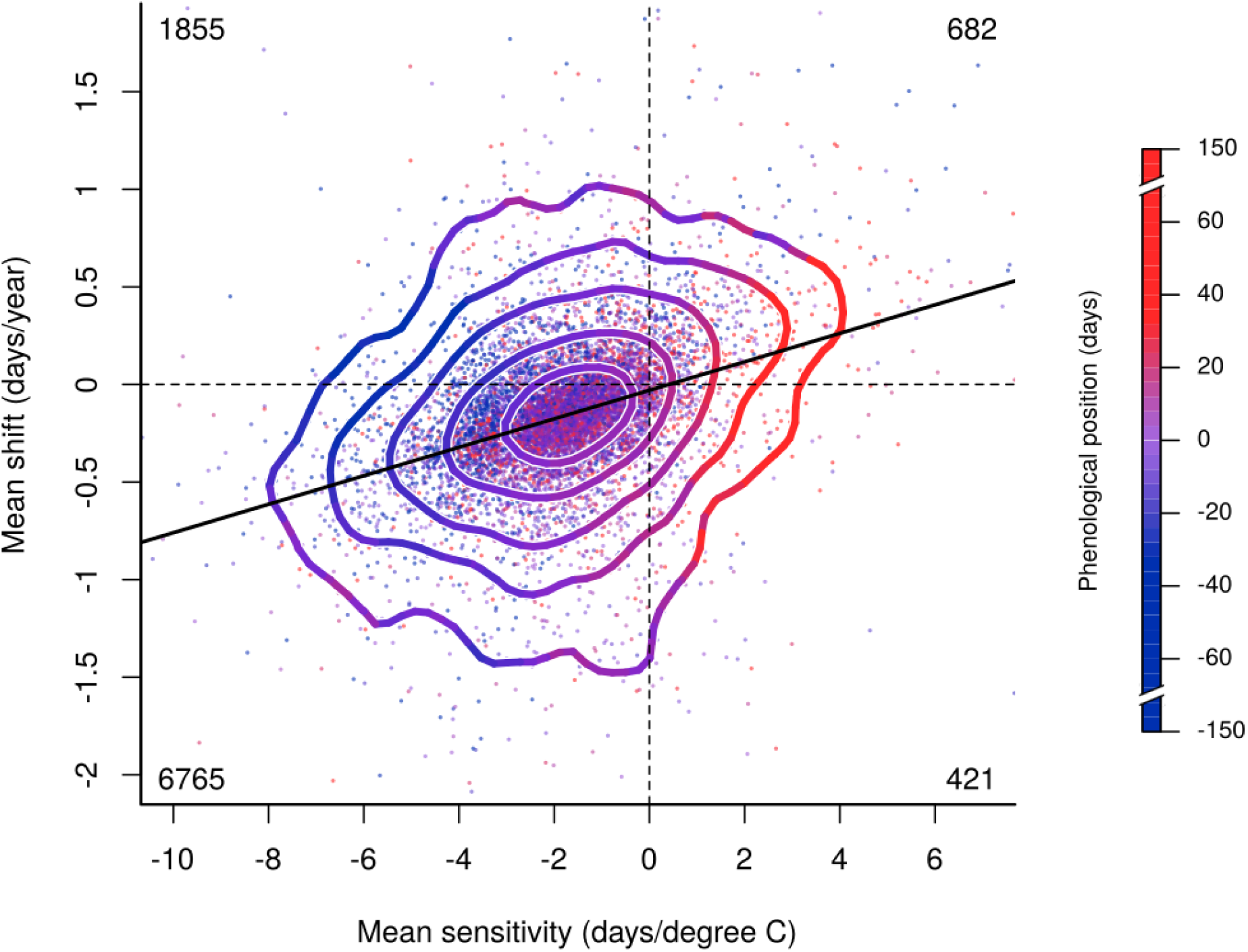
The mean sensitivity of phenology to temperature variation predicts observed shifts over time. The phenological position of species relative to others at the same sites (point and contour color; color legend on the right) is also a strong predictor of temperature sensitivity. Species whose phenophases occur on average earlier in the spring season (blue points) are more sensitive to temperature than those close to the middle (pink points) or end (red points) of spring. Most time-series exhibited both an advance in spring phenology over time and with increased temperature (bottom left), though some delayed over time but advanced with increased temperature (top left). Relatively few time-series showed a delay with increased temperature (right). Contour lines are colored by the mean phenological position of points within 0.5 mean sensitivity units and 0.25 mean shift units around the contours.

All metrics except for variance shift were predicted significantly by climatic variables and phenological position (Figure 5 top panel). Early-season species (phenological position) and those in colder regions advanced their phenology the most over time and in warmer years. Of the continuous variables, seasonality was the strongest predictor of mean sensitivity and variance sensitivity, with less seasonal areas showing the greatest effect of warm years on advancing phenology and reducing variance around that advance. Phenophase group varied in the degree of their means shifts and sensitivities, with insects advancing more than plants, and birds being the least sensitive (Figure 5 bottom panel). Phenophase groups also varied in the degree to which their phenology decreased in variance in warmer years, and variance decreased over time in leaves and flowers but not in insects and birds. The effects of all predictors were less pronounced on variance changes than they were on the corresponding mean changes. Full model coefficients and statistical results are available in Table S7.1, and are summarized in Figure 5.

**Figure 5.**
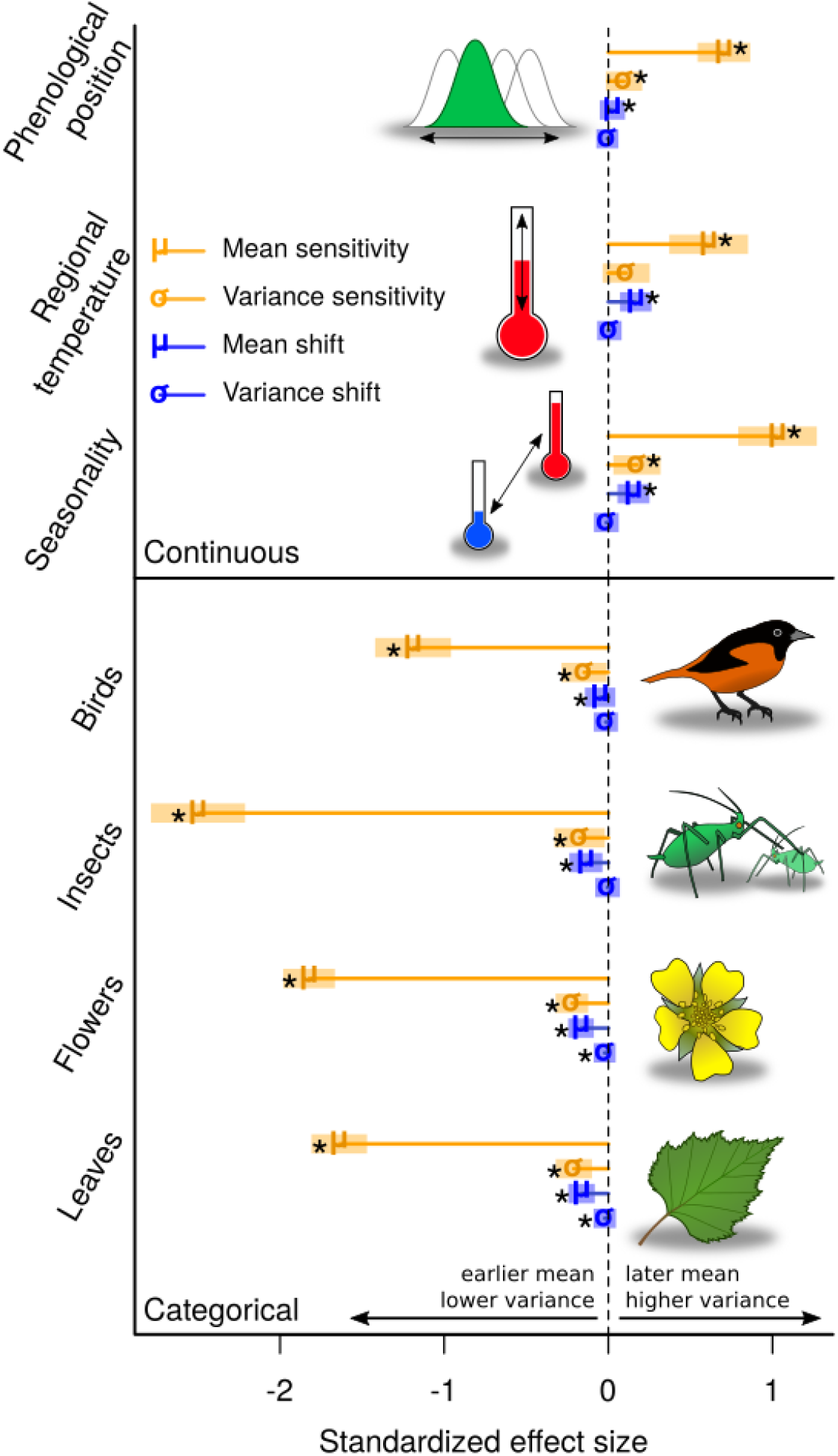
Phenological shifts and sensitivities vary by taxonomic group, regional climate, and the phenological position of species. Earlier-season species in the coldest and least seasonal areas have advanced their spring phenology and are the most sensitive to temperature variation in mean and variance (top panel). All phenophase groups advanced their mean phenology over time and in warmer years, with insects being the most sensitive (bottom panel). While variance sensitivity decreased in warmer years for all phenophase groups, variance shifted only in flowers and leaves, decreasing over time. The standardized effects of each predictor variable on the four phenological response metrics are grouped together in rows. Orange lines represent sensitivities with respect to yearly temperature variation, and blue lines represent shifts over time. Mean coefficients are represented with a μ and variance with a σ. For continuous variables, coefficients are slope parameters, and for categorical variables, coefficients are contrasts from zero. Asterisks indicate significant effects (p < 0.01), and the shaded bars represent 2×standard error around the coefficient.

Shifts in first flowering phenology and variance were not significantly predicted by the growth form of plants, height, seed mass, or SLA (Figures S4.1 – S4.4). Rates of shift in flower and leaf phenology also did not differ significantly and did not interact with growth form (Figure S4.5). Neither diet nor mass predicted phenological shifts or sensitivities in birds (Figure S5.1). Plant trait model coefficients and statistical results are available in Table S7.2, those for the phenophase model are in Table S7.3, and those for bird traits are in Table S7.4. All model results are elaborated and visualized in Supplement 3.

## Discussion

Climate change has not resulted in a uniform shift of spring phenology across all species or all parts of the world. Phenological responses have varied across trophic levels (Thackeray et al., 2016) and regional climates (Li et al., 2019; Roslin et al., 2021). Even closely related, co-occurring species can differ in their phenological responses based on their traits (Diamond et al., 2011; Konig et al., 2018) and their phenological position in the season (Cook et al., 2012; Menzel et al., 2006). While numerous factors determine rates of phenological shifts and sensitivity, some trends are general and predictable. Making predictions based on these patterns is crucial to anticipating phenological mismatches between interacting species (Renner et al., 2018) and minimizing their negative consequences on ecosystems through management (Olliff-Yang et al., 2020). Here, we confirm that even when viewed across major climatic gradients and monitoring programs in the Northern Hemisphere (Figure 2), there are consistent patterns in which species are most responsive to climate change. Moreover, we did not find evidence that climate change is making phenology inherently more variable across years. Rather, we found that the inter-annual phenological variability of plant, insect, and bird phenophases decreases in warmer years. The timings of leaf-out and flowering onset have even become modestly but significantly less variable over time, suggesting that the novel, warmer conditions presented by climate change may increase the predictability of phenology in the future.

### Predictors of phenological mean shifts and sensitivity

Some places have experienced greater changes in phenology than others based on regional climate differences. Phenology has advanced most rapidly and is most sensitive to inter-annual temperature variation in colder regions (Figure 5 top panel). Growing seasons are shorter in areas with colder climates, so plants, insects, and birds must time their activity more precisely to occur within favorable abiotic conditions (Pau et al., 2011; Roslin et al., 2021). This greater sensitivity in the colder regions of plant species distributions may lead to more connectivity and gene flow across climate gradients (Prevéy et al., 2017). In contrast to this pattern, phenology has shifted the least, is the least sensitive, and is the least inter-annually variable in the most seasonal areas after accounting for differences in the regional mean climate. In areas where temperatures change relatively little between winter and summer, species may evolve to be more sensitive to smaller inter-annual temperature differences and to more precisely track them. Viewed another way, this may simply be a product of the proportionality of phenological advance relative to temperature changes: 1°C of additional warming will have a larger relative effect on phenology in areas where the difference between winter and summer is just 10°C than in areas where the difference is 50°C.

Inherently different physiology and life-histories among taxonomic groups also determine phenological sensitivity and shifts. While all phenophase groups advanced their phenology in warmer years and over time, some were more sensitive than others (Figure 3a), with insects being the most sensitive to inter-annual temperature variation. This difference supports previous work that has found insect phenology to be more sensitive than that of plants and perhaps birds (Cohen et al., 2018; Thackeray et al., 2016). We emphasize, however, that we present less data on insects and birds than on plant phenophases, and that observed phenological trends of migratory birds are largely dependent on conditions in the region from which they are traveling, while insects are dependent on the environment at or closer to where they are active. Despite these differences among groups, there are also commonalities, with early-season species being the most sensitive and shifting their phenology earlier (Figure 4), and phenology becoming less variable in warmer years among all groups (Figure 5, bottom panel). This finding, together with a decrease in variance sensitivity in earlier species (Figure 5, top panel), supports the idea that species on the edge of their environmental tolerance have evolved to more precisely track tolerable conditions because plasticity is most consequential on the margins of climatic niches (Duputié et al., 2015). The consequences of premature leaf-out, for example, are greater in the early season (Inouye, 2008; Pardee et al., 2018) and species have evolved mechanisms such as chilling requirements to prevent leafing-out too early (Vitasse et al., 2014).

### Mechanisms of changing phenological variability

There are many plausible, potentially conflicting mechanisms that may have led to our observation of reduced phenological variability in warmer years and over time in leaf-out and flower timing (Figure 5, bottom panel). First and perhaps most obviously, inter-annual phenological variation is tied to inter-annual variation in climate. If spring temperatures become more variable between years, spring phenology should also become more variable. But the expectation of more climatic variability may not be borne out broadly in observations, as changes in inter-annual climate variance have been geographically heterogeneous (Liu et al., 2020), and we found an overall slight reduction in inter-annual temperature variance at the sites represented in this study (Figure S6.5). The observation that climate change leads to more extreme weather events within seasons does not necessarily mean that we should expect more extreme years when the overall, mean trends of climate change are accounted for (Ummenhofer et al., 2017). Another possible explanation, because population size can affect phenological first-observation dates (Miller-Rushing et al., 2008), is that decreasing populations result in later and more variable appearance observations (Figure S6.3). Many insect populations are declining (Hallmann et al., 2017), and birds that depend on insects are following suite (Bowler et al., 2019), so these declines might have counteracted decreasing inter-annual climate variability to produce no observable variance shift in insects and birds over time (Figure 5).

Beyond technical considerations, phenological mean shifts themselves may affect variance shifts. When spring phenology shifts earlier in the season, species become subject to novel environmental constraints that may affect the shape of their phenological distributions and consequently their inter-annual variance. Because the early season presents adverse conditions such as frost (Inouye, 2008; Pardee et al., 2018) and may increase the dominance of constraining phenological cues such as photoperiod (Meng et al., 2021), species’ phenological onset can become more abrupt and less variable year-to-year (Figure S6.4). These sorts of constraints may also be evidenced by nonlinear phenological responses to temperature, with species being more sensitive in colder years (Fu et al., 2015; Mulder et al., 2017), and are a plausible explanation for the observed reduction in variance sensitivity across all groups (Figure 5). In fact, non-linear responses can present themselves with reductions in variance when data are fit with linear models (Wolkovich et al., 2021), though we still observed overall negative variance shifts and sensitivity after accounting for potential non-linearity (Supplement 2). In contrast to observed patterns of decreasing variation over years, phenological variation among individuals within seasons has been shown to increase in warm years (Zohner et al., 2018), suggesting that the *intra*-annual variability of phenology does not directly translate into its *inter*-annual variability. Due to these multiple, potentially counteracting mechanisms, it is unclear how climate change will affect phenological variance going forward, and further studies are needed to investigate the relative strengths of the above mechanisms.

### Consequences of changing variability

We have argued that the inter-annual variability of phenology may decrease as climate change progresses. Although to a lesser degree than phenological mean changes, we found that variability decreases in warmer years in all phenophase groups and that the variability of flowering and leaf-out timing has already decreased over time (Figure 5, bottom panel). These findings suggest that, at least in terms of inter-annual phenological patterns, climate change may not result in ever increasing disorder, but rather a new, shifted order of species activity. Because the amount of variance around mean phenological shifts or sensitivities determines the predictability of that mean, we might expect more predictable phenological patterns in the future. From an ecological standpoint, more predictable phenologies could have wide-ranging consequences for interacting species. If leaf-out and flowering phenology continue to become more predictable over time, competitors and mutualists such as herbivores and pollinators may be more able to match their activity to the availability of plant resources if they track similar climatic cues. On the other hand, less inter-annual variability in resource phenology may be detrimental to consumers that are not able to track resource phenology due to developmental constraints or simply lack of reliable cues. If phenology continues to become less variable over time, fundamentally different adaptive strategies may be selected for in some species, with plastic responses becoming more advantageous than bet-hedging in more predictable environments (Botero et al., 2015). Changes in voltinism and bet-hedging strategies are already expected to occur as a result of warmer mean temperatures due to climate change (Dyck et al., 2015; Forrest et al., 2019), and variance shifts may accelerate these changes. If shifts in phenological means and variance due to climate change outpace species’ plasticity or abilities to adapt their strategies for phenological synchrony (Richardson et al., 2017), active management such as diversifying genotypes by relocating individuals (Olliff-Yang et al., 2020) may be needed to avoid the worst consequences of phenological mismatches for ecosystem services. A potentially positive result of increasingly predictable phenology could be in aiding climate change adaptation strategies by improving the precision of ecological forecast models that are designed to inform resource management (Enquist et al., 2014).

## Conclusion

Climate change affects phenology not only by shifting mean event dates due to species’ sensitivity to temperature, but also by changing the variation around those means. In this study, we found that the inter-annual variation of leaf-out, start of flowering, insect first-occurrence, and bird first-arrival phenology decreased in warmer years and has decreased over time in some groups. Further, regional climate and the phenological position of species help predict shifts of phenological mean shifts, sensitivity of means to temperature, and the sensitivity of variance to temperature. Our findings suggest that climate change will not necessarily lead to increasingly unpredictable inter-annual phenology, but may result in more predictable phenological patterns. Multiple conflicting factors including inter-annual climate variability, population size, environmental constraints, non-linear temperature responses, and changing intra-annual variability may be shaping phenological variance. We hope that testing the prevalence and relative importance of these mechanisms will provide avenues for further investigation and that future studies of phenology will examine changes both in inter-annual means and variances.

## Acknowledgments

M.S. was funded by the NSF Graduate Research Fellowship under Grant No. 1745048. WDP and the Pearse Lab are funded by National Science Foundation grants ABI-1759965, and EF-1802605, and UKRI/NERC NE/V009710/1. The Rothamsted Insect Survey, a National Capability, is funded by the Biotechnology and Biological Sciences Research Council under the Core Capability Grant BBS/E/C/000J0200. Funding information for the data sources are provided in the references under *Phenology data*. We thank Amanda Gallinat, Jacob Stuivenvolt-Allen, Elizabeth Wolkovich, and Jonathan Davies for providing helpful feedback on the analysis and an earlier version of the paper.

## Author Contributions

M.S. and W.D.P designed research and performed analysis; J.R.B., E.R.E, B.D.I., H.K., S.D.L, T.L.E., R.B.P., and B.T. provided data, data collection methods, and systems expertise; M.S. wrote the paper with feedback and suggestions from all authors.

## References

Abatzoglou, J. T., Dobrowski, S. Z., Parks, S. A., & Hegewisch, K. C. (2018). TerraClimate, a high-resolution global dataset of monthly climate and climatic water balance from 1958-2015. Scientific Data, 5, 1–12. https://doi.org/10.1038/sdata.2017.191

Allen, J. M., Terres, M. A., Katsuki, T., Iwamoto, K., Kobori, H., Higuchi, H., Primack, R. B., Wilson, A. M., Gelfand, A., & Silander, J. R. (2014). Modeling daily flowering probabilities: expected impact of climate change on Japanese cherry phenology. Global Change Biology, 20(4), 1251– 1263. https://doi.org/10.1111/gcb.12364

Bartomeus, I., Ascher, J. S., Wagner, D., Danforth, B. N., Colla, S., Kornbluth, S., & Winfree, R. (2011). Climate-associated phenological advances in bee pollinators and bee-pollinated plants. Proceedings of the National Academy of Sciences, 108(51), 20645–20649. https://doi.org/10.1073/pnas.1115559108

Basler, D. (2016). Evaluating phenological models for the prediction of leaf-out dates in six temperate tree species across central Europe. Agricultural and Forest Meteorology, 217, 10–21. https://doi.org/10.1016/j.agrformet.2015.11.007

Bates, D., Mächler, M., Bolker, B. M., & Walker, S. C. (2020). Linear Mixed-Effects Models using “Eigen” and S4 (1.1-26). R package.

Bell, J. R., Alderson, L., Izera, D., Kruger, T., Parker, S., Pickup, J., Shortall, C. R., Taylor, M. S., Verrier, P., & Harrington, R. (2015). Long-term phenological trends, species accumulation rates, aphid traits and climate: Five decades of change in migrating aphids. Journal of Animal Ecology, 84(1), 21–34. https://doi.org/10.1111/1365-2656.12282

Botero, C. A., Weissing, F. J., Wright, J., & Rubenstein, D. R. (2015). Evolutionary tipping points in the capacity to adapt to environmental change. Proceedings of the National Academy of Sciences of the United States of America, 112(1), 184–189. https://doi.org/10.1073/pnas.1408589111

Bowler, D. E., Heldbjerg, H., Fox, A. D., de Jong, M., & Böhning-Gaese, K. (2019). Long-term declines of European insectivorous bird populations and potential causes. Conservation Biology, 33(5), 1120–1130. https://doi.org/10.1111/cobi.13307

Breheny, P., & Burchett, W. (2015). Visualization of regression models using visreg, R package version 2.2-0. 1–15.

Burrows, M. T., Schoeman, D. S., Buckley, L. B., Moore, P., Poloczanska, E. S., Brander, K. M., Brown, C., Bruno, J. F., Duarte, C. M., Halpern, B. S., Holding, J., Kappel, C. V., Kiessling, W., O’Connor, M. I., Pandolfi, J. M., Parmesan, C., Schwing, F. B., Sydeman, W. J., & Richardson, A. J. (2011). The pace of shifting climate in marine and terrestrial ecosystems. Science, 334(6056), 652–655. https://doi.org/10.1126/science.1210288

Butler, C. J. (2003). The disproportionate effect of global warming on the arrival dates of short-distance migratory birds in North America. Ibis, 145(3), 484–495. https://doi.org/10.1046/j.1474-919X.2003.00193.x

CaraDonna, P. J., Iler, A. M., & Inouye, D. W. (2014). Shifts in flowering phenology reshape a subalpine plant community. Proceedings of the National Academy of Sciences, 111(13), 4916– 4921. https://doi.org/10.1073/pnas.1323073111

Cohen, J. M., Lajeunesse, M. J., & Rohr, J. R. (2018). A global synthesis of animal phenological responses to climate change. Nature Climate Change, 8(3), 224–228. https://doi.org/10.1038/s41558-018-0067-3

Cook, B. I., Wolkovich, E. M., Davies, T. J., Ault, T. R., Betancourt, J. L., Allen, J. M., Bolmgren, K., Cleland, E. E., Crimmins, T. M., Kraft, N. J. B., Lancaster, L. T., Mazer, S. J., McCabe, G. J., McGill, B. J., Parmesan, C., Pau, S., Regetz, J., Salamin, N., Schwartz, M. D., & Travers, S. E. (2012). Sensitivity of Spring Phenology to Warming Across Temporal and Spatial Climate Gradients in Two Independent Databases. Ecosystems, 15(8), 1283–1294. https://doi.org/10.1007/s10021-012-9584-5

Davidian, M., & Carroll, R. J. (1987). Variance function estimation. Journal of the American Statistical Association, 82(400), 1079–1091. https://doi.org/10.1080/01621459.1987.10478543

de Keyzer, C. W., Rafferty, N. E., Inouye, D. W., & Thomson, J. D. (2017). Confounding effects of spatial variation on shifts in phenology. Global Change Biology, 23(5), 1783–1791. https://doi.org/10.1111/gcb.13472

Diamond, S. E., Frame, A. M., Martin, R. A., & Buckley, L. B. (2011). Species’ traits predict phenological responses to climate change in butterflies. Ecology, 92(5), 1005–1012. https://doi.org/10.1890/10-1594.1

Díaz, S., Hodgson, J. G., Thompson, K., Cabido, M., Cornelissen, J. H. C., Jalili, A., Montserrat-Martí, G., Grime, J. P., Zarrinkamar, F., Asri, Y., Band, S. R., Basconcelo, S., Castro-Díez, P., Funes, G., Hamzehee, B., Khoshnevi, M., Pérez-Harguindeguy, N., Pérez-Rontomé, M. C., Shirvany, A., … Zak, M. R. (2004). The plant traits that drive ecosystems: Evidence from three continents. Journal of Vegetation Science, 15(3), 295. https://doi.org/10.1658/1100-9233(2004)015[0295:tpttde]2.0.co;2

Doi, H., & Katano, I. (2008). Phenological timings of leaf budburst with climate change in Japan. Agricultural and Forest Meteorology, 148(3), 512–516. https://doi.org/10.1016/j.agrformet.2007.10.002

Dowle, M., & Srinivasan, A. (2021). Extension of ‘data.frame’ (1.14.0). R package.

Duputié, A., Rutschmann, A., Ronce, O., & Chuine, I. (2015). Phenological plasticity will not help all species adapt to climate change. Global Change Biology, 21(8), 3062–3073. https://doi.org/10.1111/gcb.12914

Dyck, H. Van, Bonte, D., Puls, R., Gotthard, K., & Maes, D. (2015). The lost generation hypothesis : could climate change drive ectotherms into a developmental trap ? Oikos, 124, 54–61. https://doi.org/10.1111/oik.02066

Enquist, C. A. F., Kellermann, J. L., Gerst, K. L., & Miller-Rushing, A. J. (2014). Phenology research for natural resource management in the United States. International Journal of Biometeorology, 58(4), 579–589. https://doi.org/10.1007/s00484-013-0772-6

Ettinger, A. K., Gee, S., & Wolkovich, E. M. (2018). Phenological sequences: how early-season events define those that follow. American Journal of Botany, 105(10), 1771–1780. https://doi.org/10.1002/ajb2.1174

Forrest, J. R. K., Cross, R., & Caradonna, P. J. (2019). Two-year bee, or not two-year bee? How voltinism is affected by temperature and season length in a high-elevation solitary bee. The American Naturalist, 193(4), 560–574. https://doi.org/10.1086/701826

Fraley, C., Raftery, A. E., Murphy, T. B., & Scrucca, L. (2012). MCLUST Version 4 for R: Normal Mixture Modeling for Model-Based Clustering, Classification, and Density Estimation.

Fu, Y. H., Zhao, H., Piao, S., Peaucelle, M., Peng, S., Zhou, G., Ciais, P., Huang, M., Menzel, A., Peñuelas, J., Song, Y., Vitasse, Y., Zeng, Z., & Janssens, I. A. (2015). Declining global warming effects on the phenology of spring leaf unfolding. Nature, 526(7571), 104–107. https://doi.org/10.1038/nature15402

Gelman, A. (2008). Scaling regression inputs by dividing by two standard deviations. September 2007, 2865–2873. https://doi.org/10.1002/sim

Hallmann, C. A., Sorg, M., Jongejans, E., Siepel, H., Hofland, N., Schwan, H., Stenmans, W., Müller, A., Sumser, H., Hörren, T., Goulson, D., & De Kroon, H. (2017). More than 75 percent decline over 27 years in total flying insect biomass in protected areas. PLoS ONE, 12(10). https://doi.org/10.1371/journal.pone.0185809

Ibáñez, I., Primack, R. B., Miller-Rushing, A. J., Ellwood, E., Higuchi, H., Lee, S. D., Kobori, H., & Silander, J. A. (2010). Forecasting phenology under global warming. Philosophical Transactions of the Royal Society B: Biological Sciences, 365(1555), 3247–3260. https://doi.org/10.1098/rstb.2010.0120

Iler, A. M., Høye, T. T., Inouye, D. W., & Schmidt, N. M. (2013). Nonlinear flowering responses to climate: Are species approaching their limits of phenological change? Philosophical Transactions of the Royal Society B: Biological Sciences, 368(1624), 13–16. https://doi.org/10.1098/rstb.2012.0489

Inouye, B. D., Ehrlén, J., & Underwood, N. (2019). Phenology as a process rather than an event: from individual reaction norms to community metrics. Ecological Monographs, 89(2), 1–15. https://doi.org/10.1002/ecm.1352

Inouye, D. W. (2008). Effects of Climate Change on Phenology, Frost Damage, and Floral Abundance of Montane Wildflowers. Ecology, 89(2), 353–362.

Keenan, T. F., Richardson, A. D., & Hufkens, K. (2020). On quantifying the apparent temperature sensitivity of plant phenology. New Phytologist, 225(2), 1033–1040. https://doi.org/10.1111/nph.16114

Kim, M., Lee, S., Lee, H., & Lee, S. (2021). Phenological response in the trophic levels to climate change in Korea. International Journal of Environmental Research and Public Health, 18(3), 1– 12. https://doi.org/10.3390/ijerph18031086

Koenker, R. (2021). Quantile regression. https://doi.org/10.1017/CBO9780511754098

Koenker, R., & Hallock, K. F. (2001). Quantile regression. Journal of Economic Perspectives, 15(4), 143–156. https://doi.org/10.1038/s41592-019-0406-y

Konig, P., Tautenhahn, S., Cornelissen, J. H. C., Christine, R., Kattge, J., & Gerhard, B. (2018). Advances in flowering phenology across the Northern Hemisphere are explained by functional traits. June 2016, 310–321. https://doi.org/10.1111/geb.12696

Kuznetsova, A., Brockhoff, P. B., & Christensen, R. H. B. (2017). lmerTest Package: Tests in Linear Mixed Effects Models. Journal of Statistical Software, 82(13). https://doi.org/10.18637/jss.v082.i13

Leung, C., Rescan, M., Grulois, D., & Chevin, L. M. (2020). Reduced phenotypic plasticity evolves in less predictable environments. Ecology Letters, 23(11), 1664–1672. https://doi.org/10.1111/ele.13598

Li Stucky, B.J., Deck, J., Baiser, B., & Guralnick, R. P. (2019). The effect of urbanization on plant phenology depends on regional temperature. Nature Ecology and Evolution, 3(12), 1661–1667. https://doi.org/10.1038/s41559-019-1004-1

Lindestad, O., Wheat, C. W., Nylin, S., & Gotthard, K. (2019). Local adaptation of photoperiodic plasticity maintains life cycle variation within latitudes in a butterfly. Ecology, 100(1), 1–14. https://doi.org/10.1002/ecy.2550

Liu, L., & Zhang, X. (2020). Effects of temperature variability and extremes on spring phenology across the contiguous United States from 1982 to 2016. Scientific Reports, 10(1), 1–14. https://doi.org/10.1038/s41598-020-74804-4

Lloyd-Evans, T. L., & Atwood, J. L. (2004). 32 Years of changes in passerine numbers during spring and fall migrations in coastal Massachusetts. Wilson Bulletin, 116(1), 1–16. https://doi.org/10.1676/0043-5643(2004)116[0001:YOCIPN]2.0.CO;2

Maitner, B. S., Boyle, B., Casler, N., Condit, R., Donoghue, J., Durán, S. M., Guaderrama, D., Hinchliff, C. E., Jørgensen, P. M., Kraft, N. J. B., McGill, B., Merow, C., Morueta-Holme, N., Peet, R. K., Sandel, B., Schildhauer, M., Smith, S. A., Svenning, J. C., Thiers, B., … Enquist, B. J. (2018). The bien r package: A tool to access the Botanical Information and Ecology Network (BIEN) database. Methods in Ecology and Evolution, 9(2), 373–379. https://doi.org/10.1111/2041-210X.12861

Meng, L., Zhou, Y., Gu, L., Richardson, A. D., Peñuelas, J., Fu, Y., Wang, Y., Asrar, G. R., Boeck, H. J. De, Mao, J., Zhang, Y., & Wang, Z. (2021). Photoperiod decelerates the advance of spring phenology of six deciduous tree species under climate warming. Global Change Biology. https://doi.org/10.1111/gcb.15575

Menzel, A., Sparks, T. H., Estrella, N., & Roy, D. B. (2006). Altered geographic and temporal variability in phenology in response to climate change. Global Ecology and Biogeography, 15(5), 498–504. https://doi.org/10.1111/j.1466-822X.2006.00247.x

Miller-Rushing, A. J., Inouye, D. W., & Primack, R. B. (2008). How well do first flowering dates measure plant responses to climate change? The effects of population size and sampling frequency. Journal of Ecology, 96(6), 1289–1296. https://doi.org/10.1111/j.1365-2745.2008.01436.x

Miller-Rushing, A. J., Lloyd-Evans, T. L., Primack, R. B., & Satzinger, P. (2008). Bird migration times, climate change, and changing population sizes. Global Change Biology, 14(9), 1959–1972. https://doi.org/10.1111/j.1365-2486.2008.01619.x

Mulder, C. P. H., Iles, D. T., & Rockwell, R. F. (2017). Increased variance in temperature and lag effects alter phenological responses to rapid warming in a subarctic plant community. Global Change Biology, 23(2), 801–814. https://doi.org/10.1111/gcb.13386

Olliff-Yang, R. L., Gardali, T., & Ackerly, D. D. (2020). Mismatch managed? Phenological phase extension as a strategy to manage phenological asynchrony in plant–animal mutualisms. Restoration Ecology, 28(3), 498–505. https://doi.org/10.1111/rec.13130

Ovaskainen, O., Meyke, E., Lo, C., Tikhonov, G., Delgado, M. del M., Roslin, T., Gurarie, E., Abadonova, M., & Abduraimov, O. (2020). Chronicles of nature calendar, a long-term and large-scale multitaxon database on phenology. Scientific Data, 7(47), 1–11. https://doi.org/10.1038/s41597-020-0376-z

Pardee, G. L., Inouye, D. W., & Irwin, R. E. (2018). Direct and indirect effects of episodic frost on plant growth and reproduction in subalpine wildflowers. April 2017, 848–857. https://doi.org/10.1111/gcb.13865

Parmesan, C. (2007). Influences of species, latitudes and methodologies on estimates of phenological response to global warming. Global Change Biology, 13(9), 1860–1872. https://doi.org/10.1111/j.1365-2486.2007.01404.x

Parmesan, C., & Yohe, G. (2003). A globally coherent fingerprint of climate change impacts across natural systems. Nature, 421, 37–42. https://doi.org/10.1038/nature01286

Patterson, T. A., Grundel, R., Dzurisin, J. D. K., Knutson, R. L., & Hellmann, J. J. (2020). Evidence of an extreme weather induced phenological mismatch and a local extirpation of the endangered Karner blue butterfly. Conservation Science and Practice, 2(e147), 1–13. https://doi.org/10.1111/csp2.147

Pau, S., Wolkovich, E. M., Cook, B. I., Davies, T. J., Kraft, N. J. B., Bolmgren, K., Betancourt, J. L., & Cleland, E. E. (2011). Predicting phenology by integrating ecology, evolution and climate science. Global Change Biology, 17(12), 3633–3643. https://doi.org/10.1111/j.1365-2486.2011.02515.x

Pearse, W. D., Davis, C. C., Inouye, D. W., Primack, R. B., & Davies, T. J. (2017). A statistical estimator for determining the limits of contemporary and historic phenology. Nature Ecology and Evolution, 1(12), 1876–1882. https://doi.org/10.1038/s41559-017-0350-0

Pebesma, E. J. (2021). Simple Features for R (0.9-7). R package.

Prevéy, J., Vellend, M., Rüger, N., Hollister, R. D., Bjorkman, A. D., Myers-Smith, I. H., Elmendorf, S. C., Clark, K., Cooper, E. J., Elberling, B., Fosaa, A. M., Henry, G. H. R., Høye, T. T., Jónsdóttir, I.S., Klanderud, K., Lévesque, E., Mauritz, M., Molau, U., Natali, S. M., … Rixen, C. (2017). Greater temperature sensitivity of plant phenology at colder sites: implications for convergence across northern latitudes. Global Change Biology, 23(7), 2660–2671. https://doi.org/10.1111/gcb.13619

Primack, R. B. (1987). Relationships Among Flowers, Fruits, and Seeds. Annual Review of Ecology and Systematics, 18(1), 409–430. https://doi.org/10.1146/annurev.es.18.110187.002205

R Core Team. (2020). R: A language and environment for statistical computing (3.6.3). R Foundation for Statistical Computing.

Renner, S. S., & Zohner, C. M. (2018). Climate change and phenological mismatch in trophic interactions among plants, insects, and vertebrates. In Annual Review of Ecology, Evolution, and Systematics (Vol. 49, pp. 165–182). https://doi.org/10.1146/annurev-ecolsys-110617-062535

Richardson, B. A., Chaney, L., Shaw, N. L., & Still, S. M. (2017). Will phenotypic plasticity affecting flowering phenology keep pace with climate change? Global Change Biology, 23(6), 2499–2508. https://doi.org/10.1111/gcb.13532

Roslin, T., Antão, L., Hällfors, M., Meyke, E., Lo, C., Tikhonov, G., Delgado, M. del M., Gurarie, E., Abadonova, M., Abduraimov, O., Adrianova, O., Akimova, T., Akkiev, M., Ananin, A., Andreeva, E., Andriychuk, N., Antipin, M., Arzamascev, K., Babina, S., … Ovaskainen, O. (2021). Phenological shifts of abiotic events, producers and consumers across a continent. Nature Climate Change, 3. https://doi.org/10.1038/s41558-020-00967-7

Smith, R. L. (1987). Estimating Tails of Probability Distributions. The Annals of Statistics, 15(3), 1174–1207.

South, A. (2017). Rnaturalearth: world map data from natural earth (0.1). R package.

Stegman, L. S., Primack, R. B., Gallinat, A. S., Lloyd-Evans, T. L., & Ellwood, E. R. (2017). Reduced sampling frequency can still detect changes in abundance and phenology of migratory landbirds. Biological Conservation, 210(April), 107–115. https://doi.org/10.1016/j.biocon.2017.04.004

Templ, B., Koch, E., Bolmgren, K., Ungersböck, M., Paul, A., Scheifinger, H., Rutishauser, T., Busto, M., Chmielewski, F. M., Hájková, L., Hodzić, S., Kaspar, F., Pietragalla, B., Romero-Fresneda, R., Tolvanen, A., Vučetič, V., Zimmermann, K., & Zust, A. (2018). Pan European Phenological database (PEP725): a single point of access for European data. International Journal of Biometeorology, 62(6), 1109–1113. https://doi.org/10.1007/s00484-018-1512-8

Thackeray, S. J., Henrys, P. A., Hemming, D., Bell, J. R., Botham, M. S., Burthe, S., Helaouet, P., Johns, D. G., Jones, I. D., Leech, D. I., MacKay, E. B., Massimino, D., Atkinson, S., Bacon, P. J., Brereton, T. M., Carvalho, L., Clutton-Brock, T. H., Duck, C., Edwards, M., … Wanless, S. (2016). Phenological sensitivity to climate across taxa and trophic levels. Nature, 535(7611), 241– 245. https://doi.org/10.1038/nature18608

Tiusanen, M., Kankaanpää, T., Schmidt, N. M., & Roslin, T. (2020). Heated rivalries: Phenological variation modifies competition for pollinators among arctic plants. Global Change Biology, 26(11), 6313–6325. https://doi.org/10.1111/gcb.15303

Ummenhofer, C. C., & Meehl, G. A. (2017). Extreme weather and climate events with ecological relevance: A review. Philosophical Transactions of the Royal Society B: Biological Sciences, 372(1723). https://doi.org/10.1098/rstb.2016.0135

Visser, M. E., & Gienapp, P. (2019). Evolutionary and demographic consequences of phenological mismatches. Nature Ecology & Evolution, 12, 879–885. https://doi.org/10.1038/s41559-019-0880-8

Vitasse, Y., Lenz, A., & Körner, C. (2014). The interaction between freezing tolerance and phenology in temperate deciduous trees. Frontiers in Plant Science, 5(OCT), 1–12. https://doi.org/10.3389/fpls.2014.00541

Wadgymar, S. M., Ogilvie, J. E., Inouye, D. W., Weis, A. E., & Anderson, J. T. (2018). Phenological responses to multiple environmental drivers under climate change: insights from a long-term observational study and a manipulative field experiment. New Phytologist, 218(2), 517–529. https://doi.org/10.1111/nph.15029

Wilke, C. O. (2020). Streamlined Plot Theme and Plot Annotations for “ggplot2” (1.1.1). R package.

Wilman, H., J., B., J., S., C., de L. R., M., R., & W, J. (2014). EltonTraits 1.0 : Species-level foraging attributes of the world’s birds and mammals. Ecology, 95(7), 2027.

Wolkovich, E., Auerbach, J., Chamberlain, C., Buonaiuto, D., Ettinger, A., Morales-Castilla, I., & Gelman, A. (2021). A simple explanation for declining temperature sensitivity with warming. Global Change Biology, 27(20), 1–3. https://doi.org/10.1111/gcb.15746

Zohner, C. M., Mo, L., & Renner, S. S. (2018). Global warming reduces leaf-out and flowering synchrony among individuals. ELife, 7(e40214), 1–15.

