## Supplementary Materials for "Disorder or a new order: how climate change affects phenological variability"

### Supplement 1. Power analysis

We performed a power analysis in order to determine the efficacy of the residual method in identifying changes in variance. We simulated time-series data by generating 10,000 sequences of 32 years, with the phenological trends varying uniformly within 40 days around the baseline phenology occurring at 100 days-of-year. For each time-series, we added normally distributed error to each time point with a baseline standard deviation of 12 days. We then increased the error term standard deviation by ten increments between -10 and +10 days, resulting in a total error standard deviation range between 2 and 22 days and 1000 replicated per variance change increment. We then performed the residual method test on each time-series and assessed success of the method by testing whether the actual variance change fell within 2 standard errors of the predicted variance change. Actual variance change was calculated as the increase/decrease in the total error SD divided by the number of data points, and the predicted variance change was the slop of the absolute residuals. We then calculated the percentage of successes in each of the 1000 variance change replicates.

We found that the residual method was over 90% successful in correctly detecting variance changes for every variance shift scenario (Fig. S1). We investigated potential biases in the estimation of mean and variance changes, but found that our method accurately predicted both metrics one-to-one. The residual method was equally likely to over-estimate variance change as it was to under-estimate it. We conclude that the residual method of fitting a quantile regression with $\tau \approx 0.6827$ to the absolute residuals accurately predicts the change in the standard deviation of the error term under the present assumptions.

**Figure S1**. The power of the residual method for different rates of variance change. Black point indicate simulations for which the method successfully predicted the rate of variance change, and red circles represent those for which it didn't. The ideal 1:1 line is solid black, the actual performance of the residual method is dashed black, and zero variance change is highlighted by the red line.

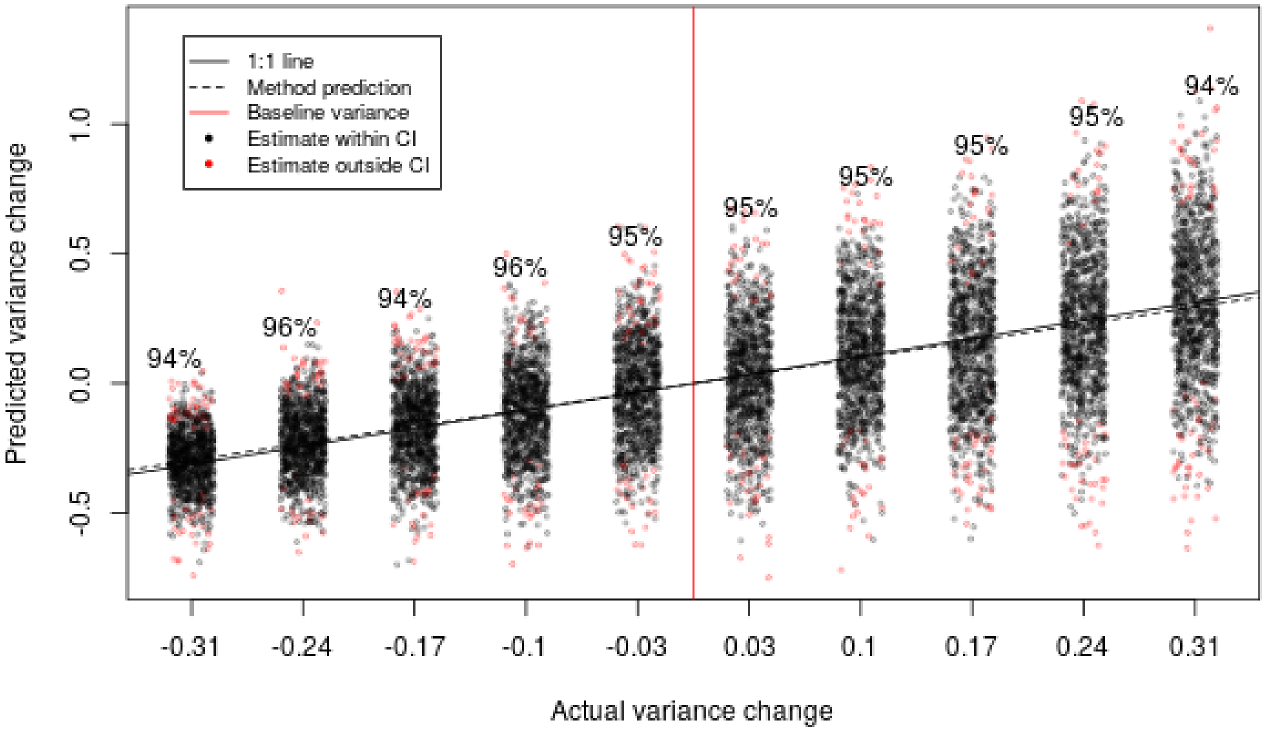

**Supplement 2. Accounting for potential non-linearity**

It is possible that if phenological responses to increasing temperature are non-linear, as is predicted by a thermal threshold model, that fitting linear models and estimating variance from their residuals might result in biased measures of variance change (Wolkovich et al., 2021). One way to account for the expected non-linearity of a thermal threshold phenological process is to log-transform the response variable when making sensitivity calculations. So, to check whether and to what extend assumptions about linear temperature responses affected our estimates of variance shifts and sensitivity, we $log_e$-transformed the response variable DOY before calculating the mean and variance metrics. We found that mean shifts and mean sensitivity were still clearly negative overall after this transformation. There were still overall trends toward negative variance sensitivity and negative variance shifts (Figure S2), but that the effects were smaller than when DOY was not transformed (Figure S3.1). This indicates that the statistical assumption of a linear relationship between inter-annual temperature and phenology may in part explain the observed reduction in variance, but variance is still observed to decrease even if the assumption is faulty.

**Figure S2**. Overall trends distributions for all phenophase groups when day-of-year is log-transformed to account for potentially non-linear relationships between phenology and temperature. The plotting range is reduced to show detail of the center of the distributions, so some points are not plotted.

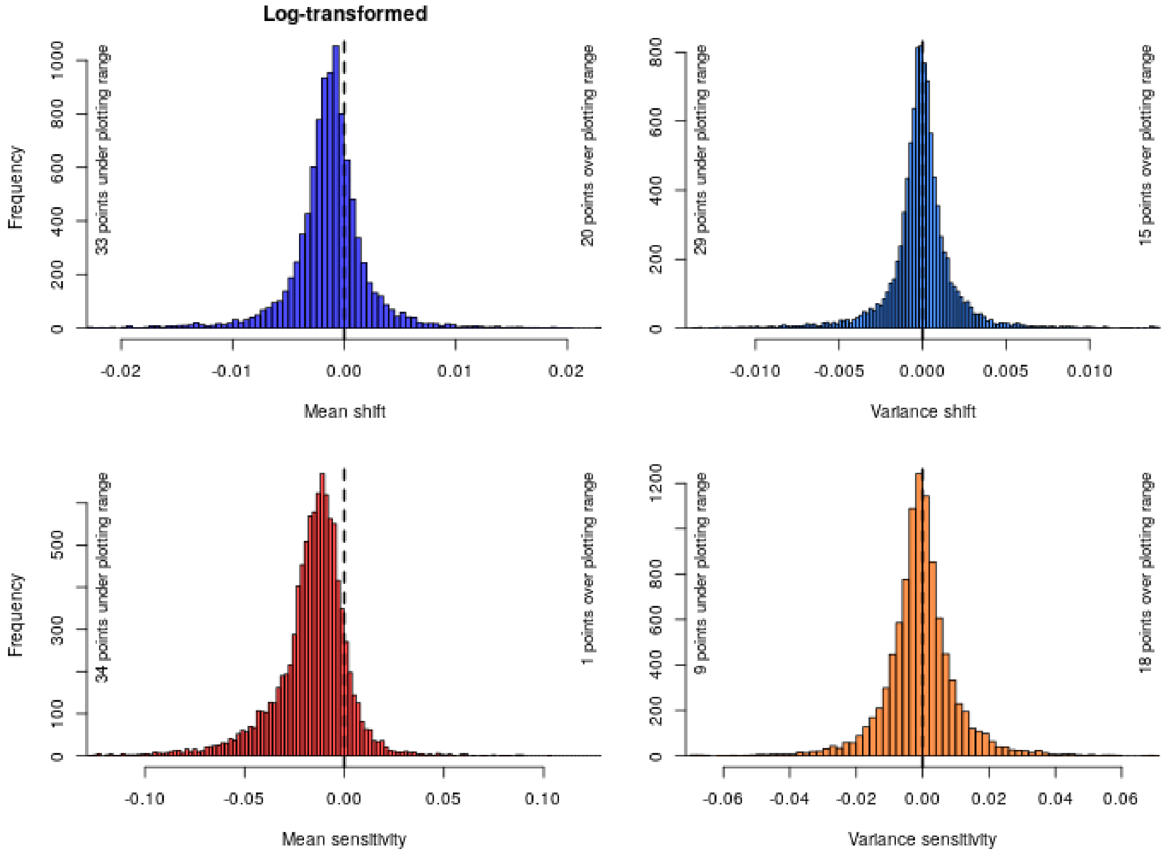

**Supplement 3. Trend visualizations**

**Figure S3.1**. Overall trends distributions for all phenophase groups. The plotting range is reduced to show detail of the center of the distributions, so some points are not plotted.

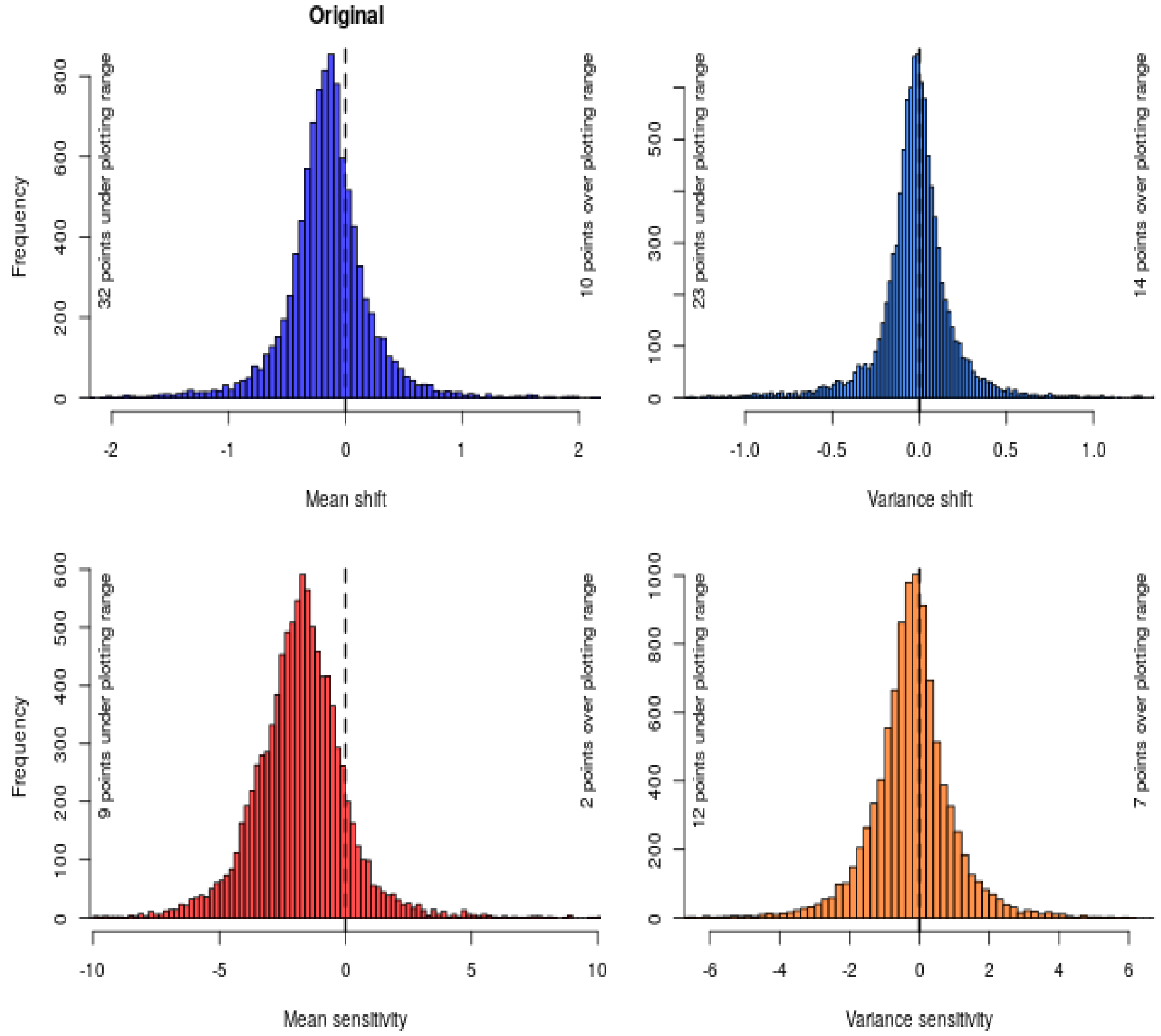

**Figure S3.2**. The drivers of phenology mean shifts over time. Lines are model predictions and points are residuals.

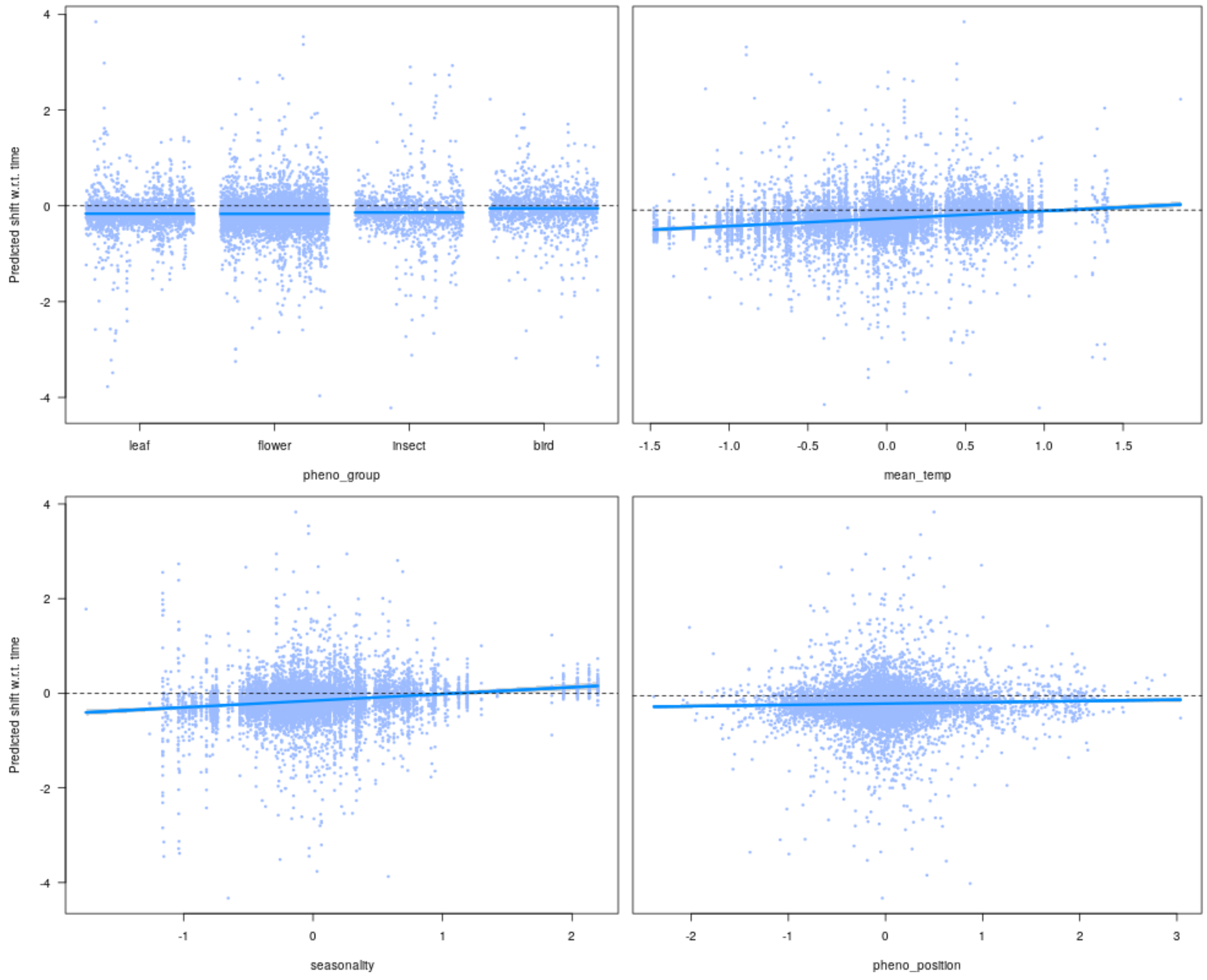

**Figure S3.3**. The drivers of the temperature sensitivity of phenological means. Lines are model predictions and points are residuals.

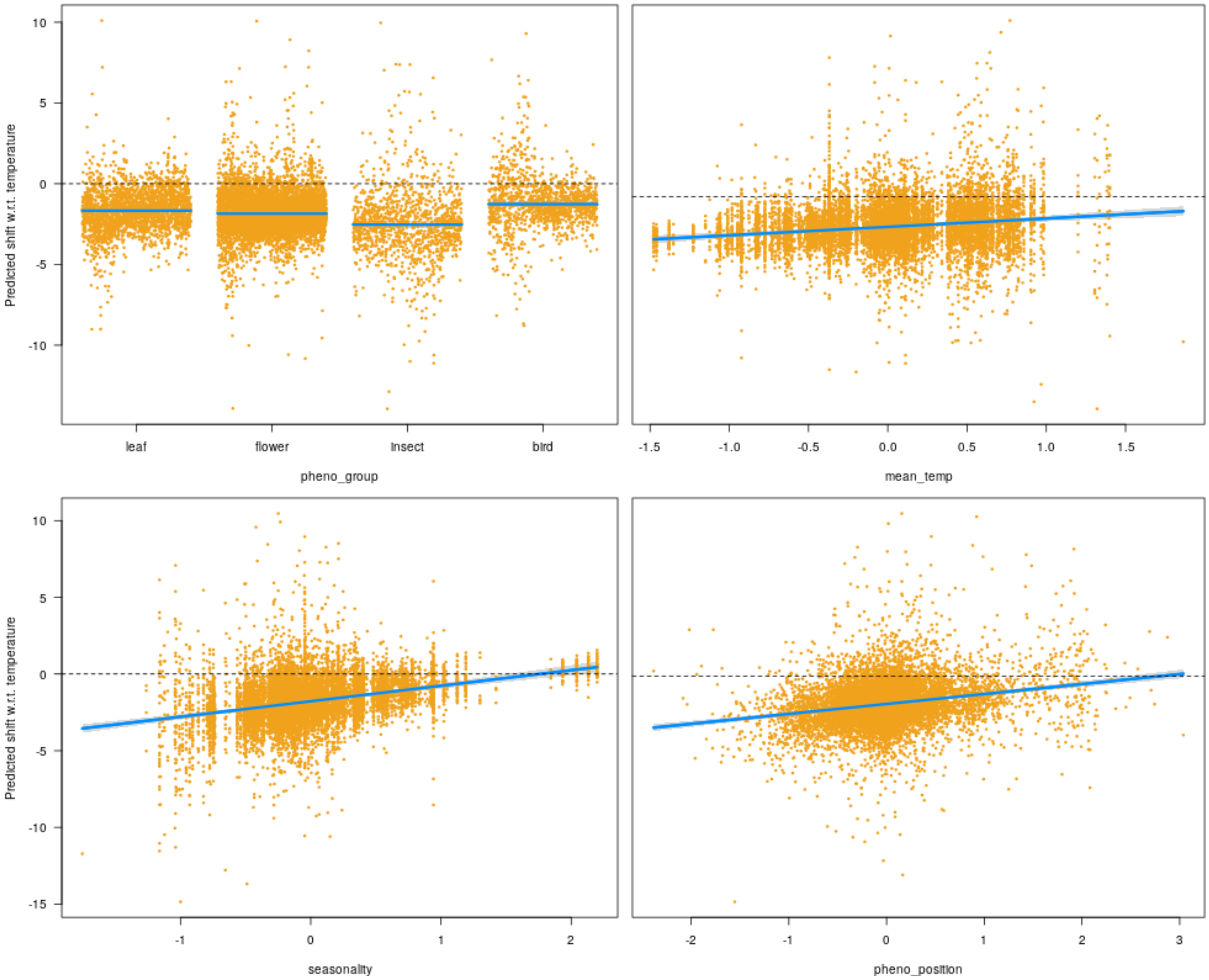

**Figure S3.4**. The drivers of variance shifts over time. Lines are model predictions and points are residuals.

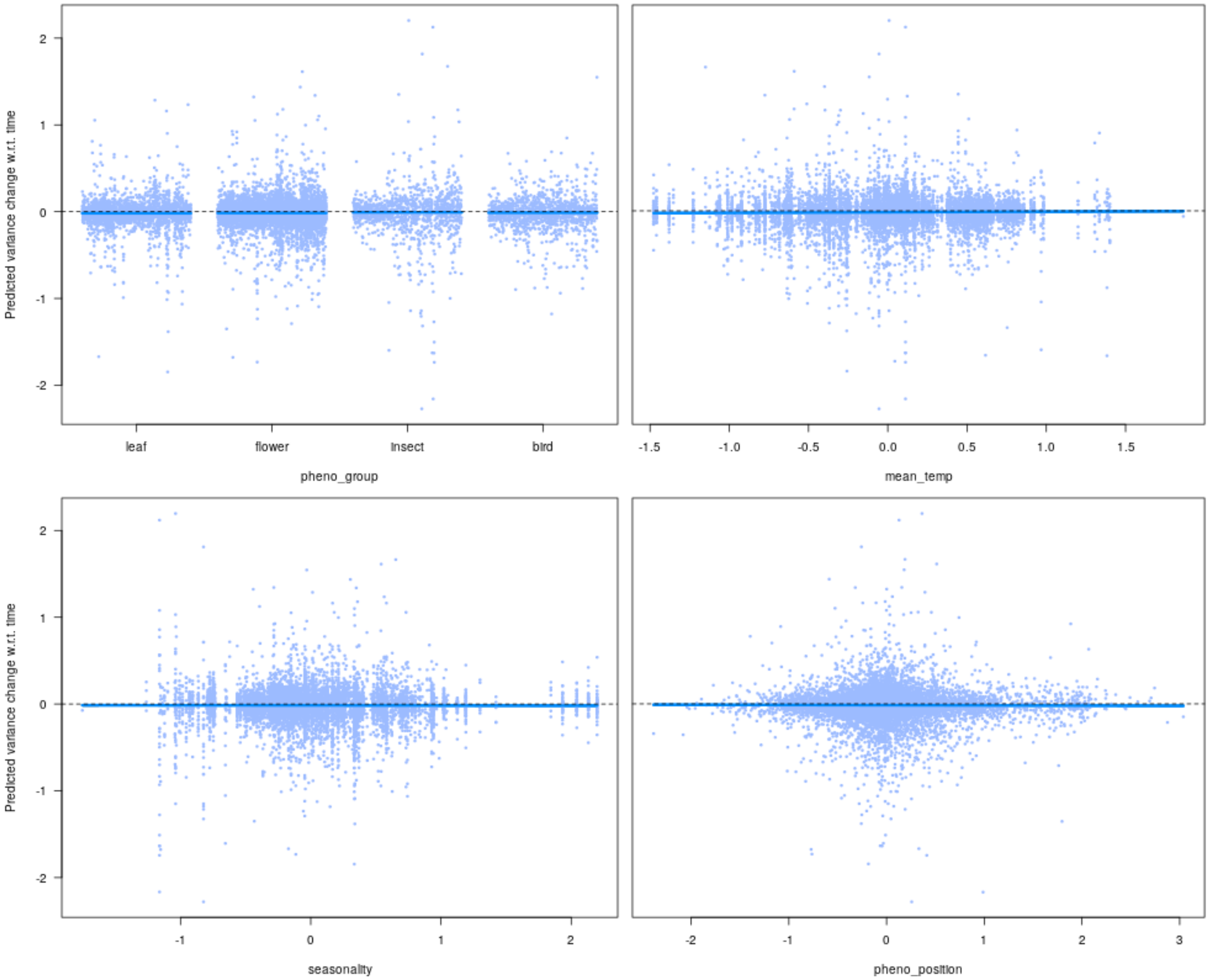

**Figure S3.5**. The drivers of variance temperature sensitivity. Lines are model predictions and points are residuals.

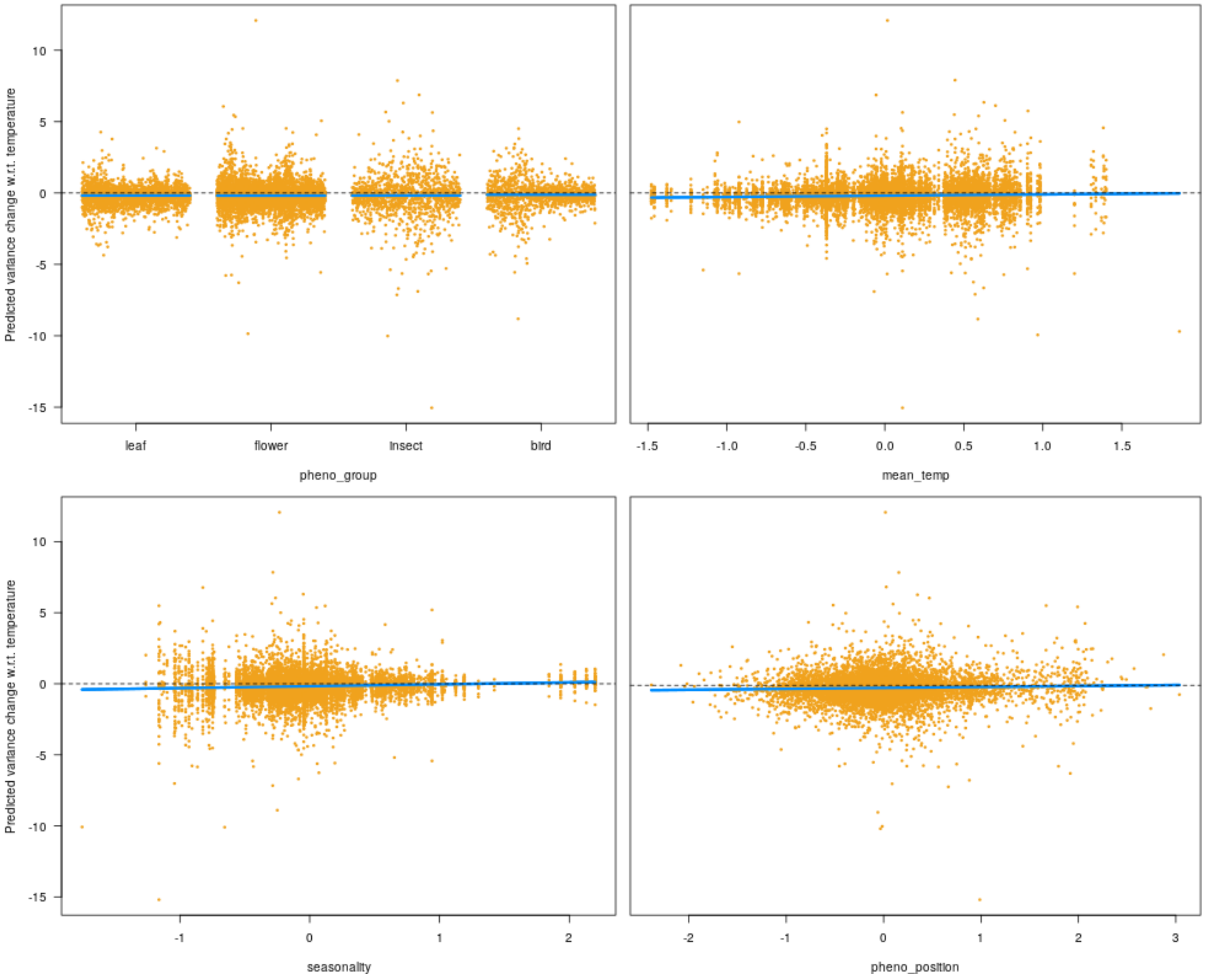

**Supplement 4. Plant trait analyses**

**Figure S4.1**. The drivers of first flowering phenology mean shifts over time. No effects were significant. Lines are model predictions and points are residuals.

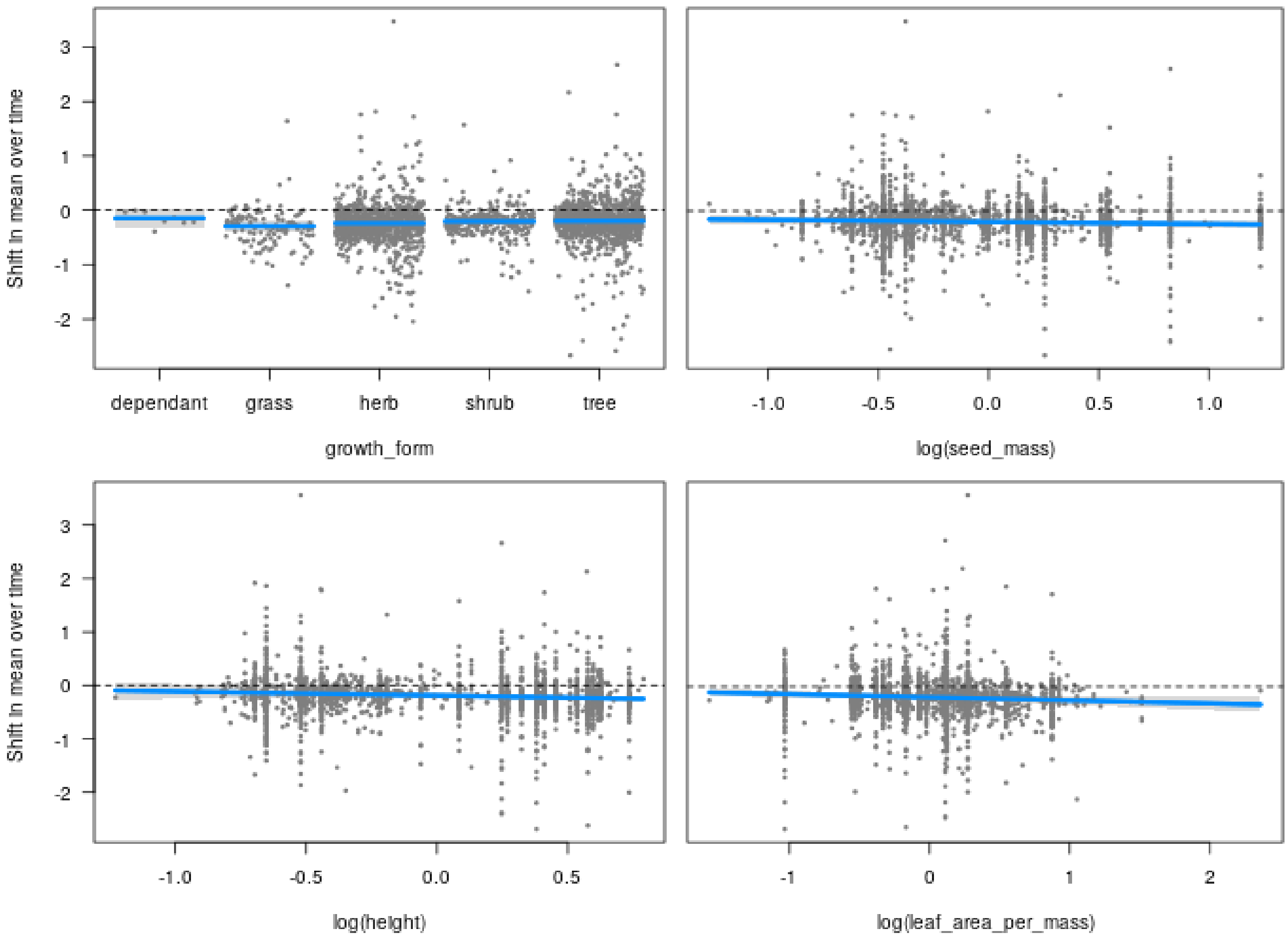

**Figure S4.2**. The drivers of first flowering mean temperature sensitivity. Dependents were significantly less sensitive than the other growth forms. Lines are model predictions and points are residuals.

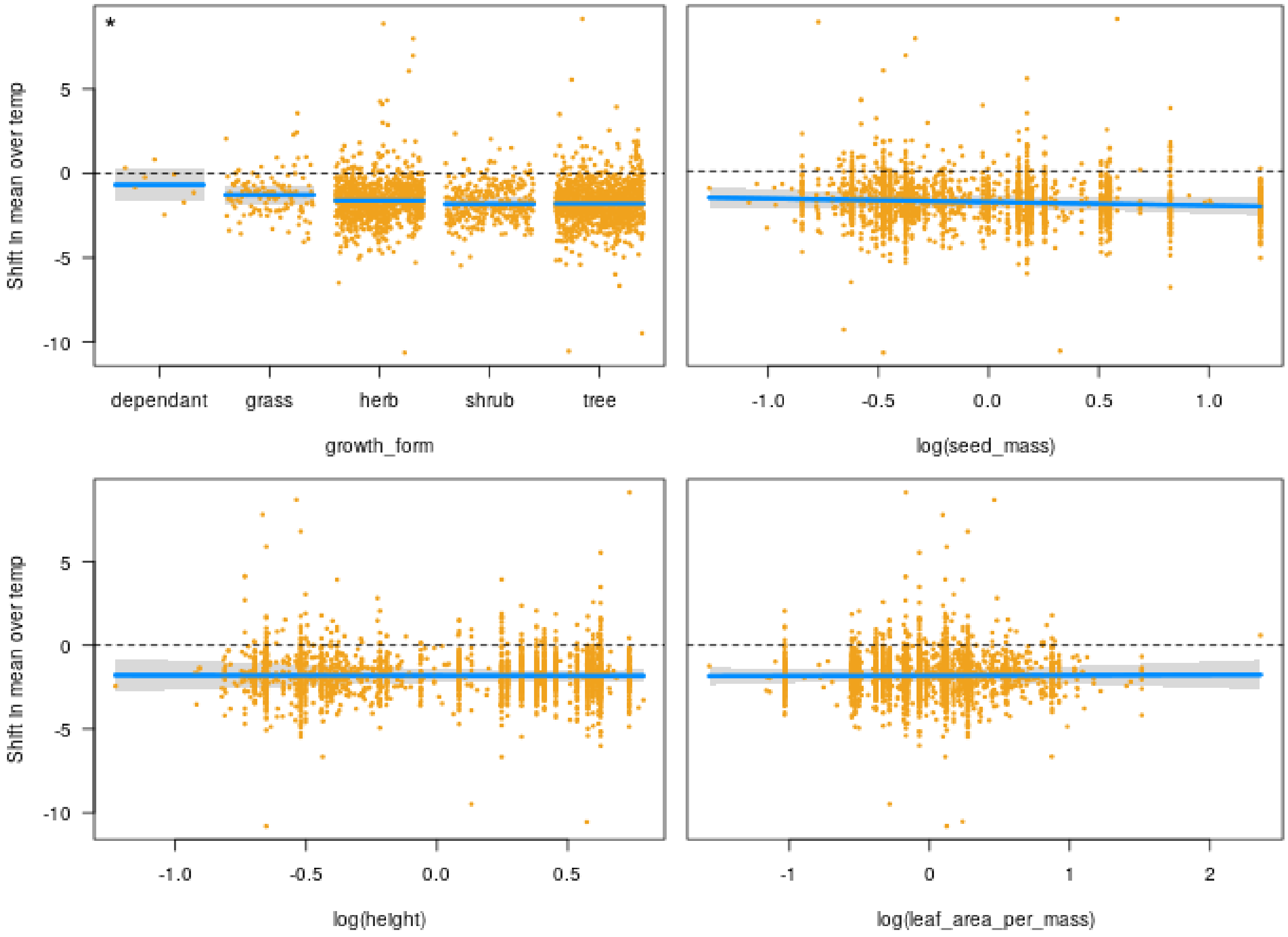

**Figure S4.3**.The drivers of variance changes in first flowering over time. No effects were significant. Lines are model predictions and points are residuals.

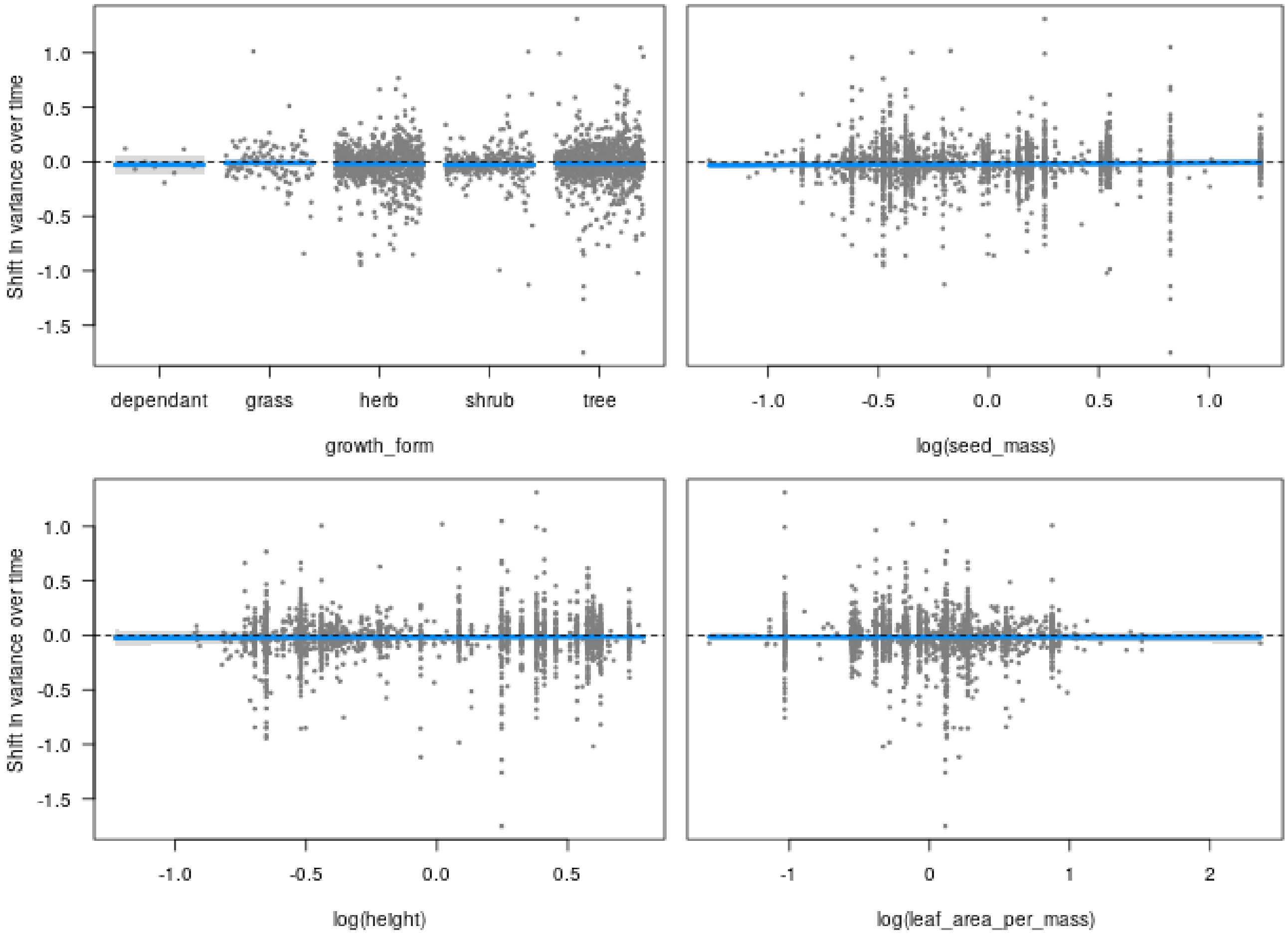

**Figure S4.4**. The drivers of the temperature sensitivity of variance changes in first flowering. No effects were significant. Lines are model predictions and points are residuals.

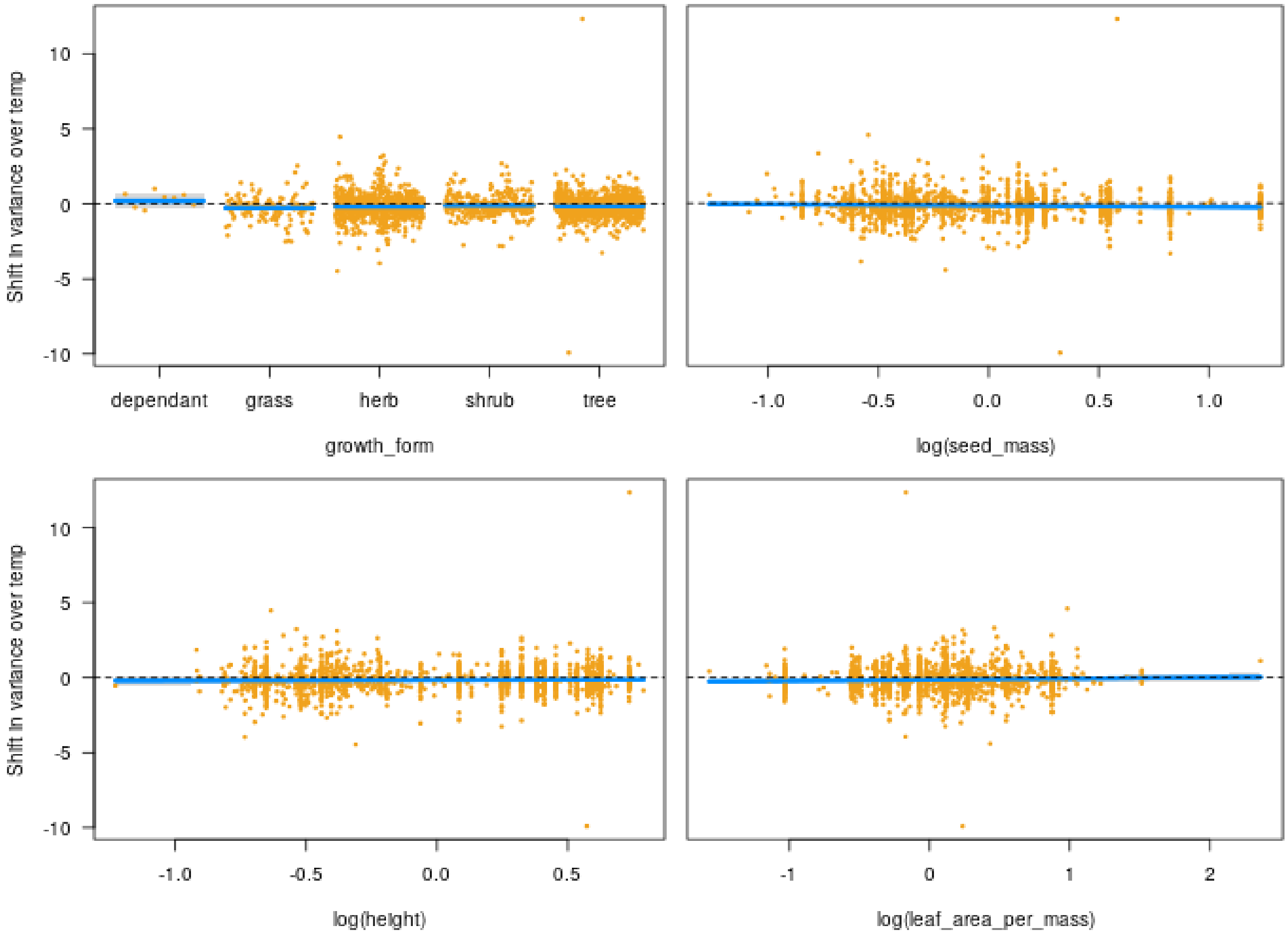

**Figure S4.5**. Neither mean phenological shifts over time, temperature sensitivity, nor changes in variance varied significantly between leaf and flower phenology. The patterns also did not significantly differ between trees, shrubs, and herbs, and there were no significant interactions between growth form and phenophase. Red lines and points are model predictions and residuals for shrubs, and blue ones are those for trees.

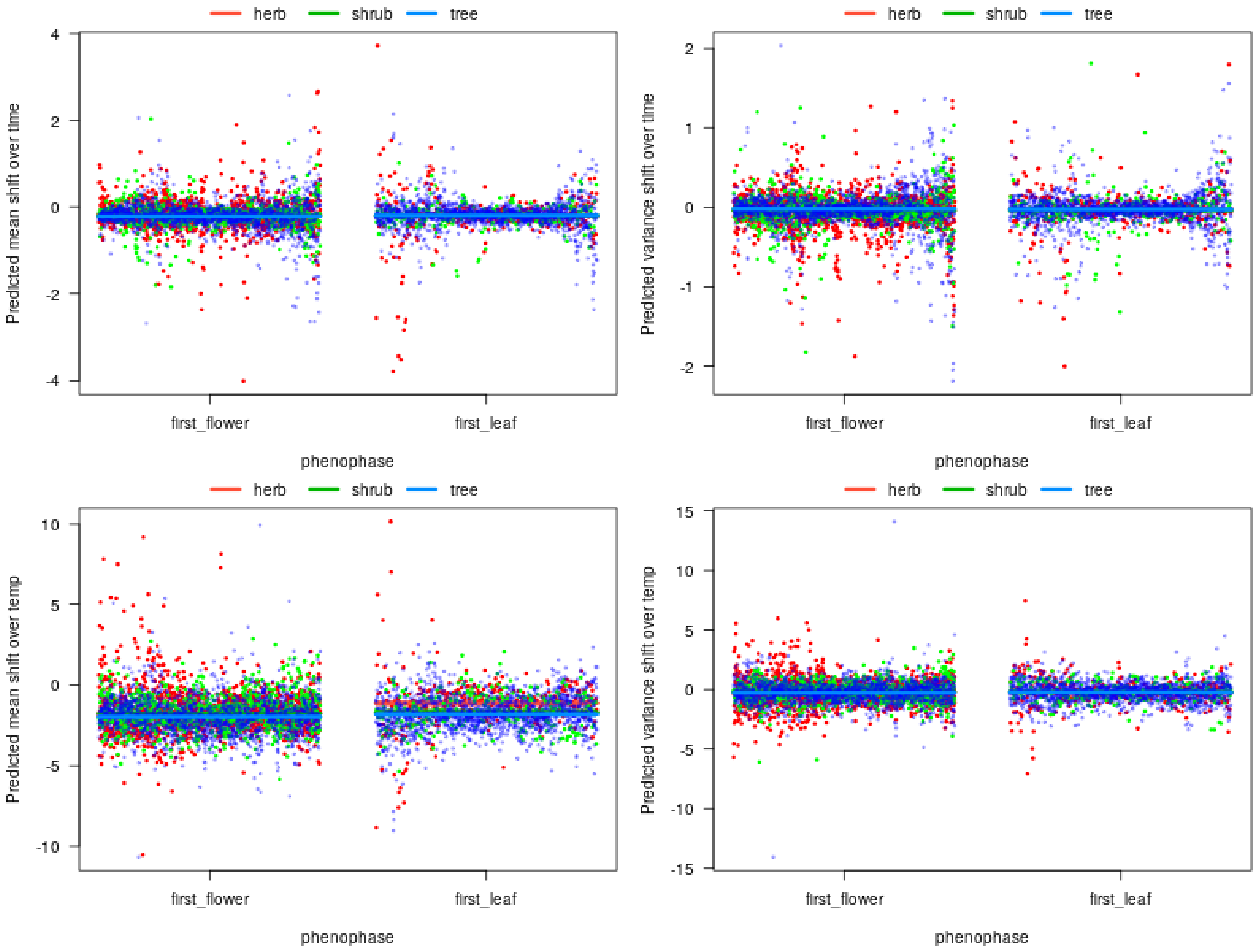

**Supplement 5. Bird trait analyses**

**Figure S5.1**. Bird diet did not significantly predict mean phenological shifts over years, temperature sensitivity, or variance changes. Lines are model predictions and points are residuals.

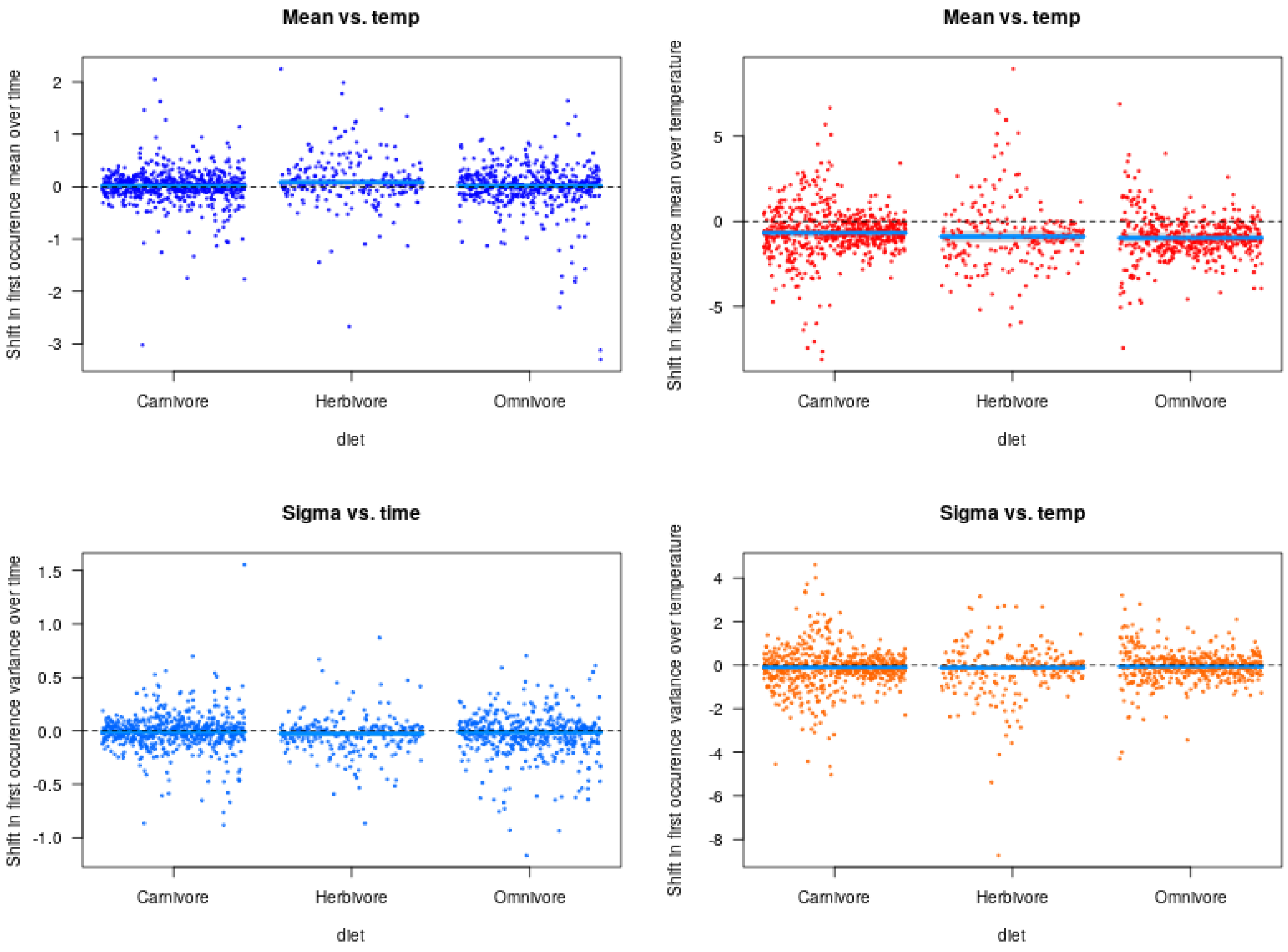

### Supplement 6. Additional analyses

**Figure S6.1**. Sites within datasets were regionally clustered and spanned different segments of the seasonality gradient.

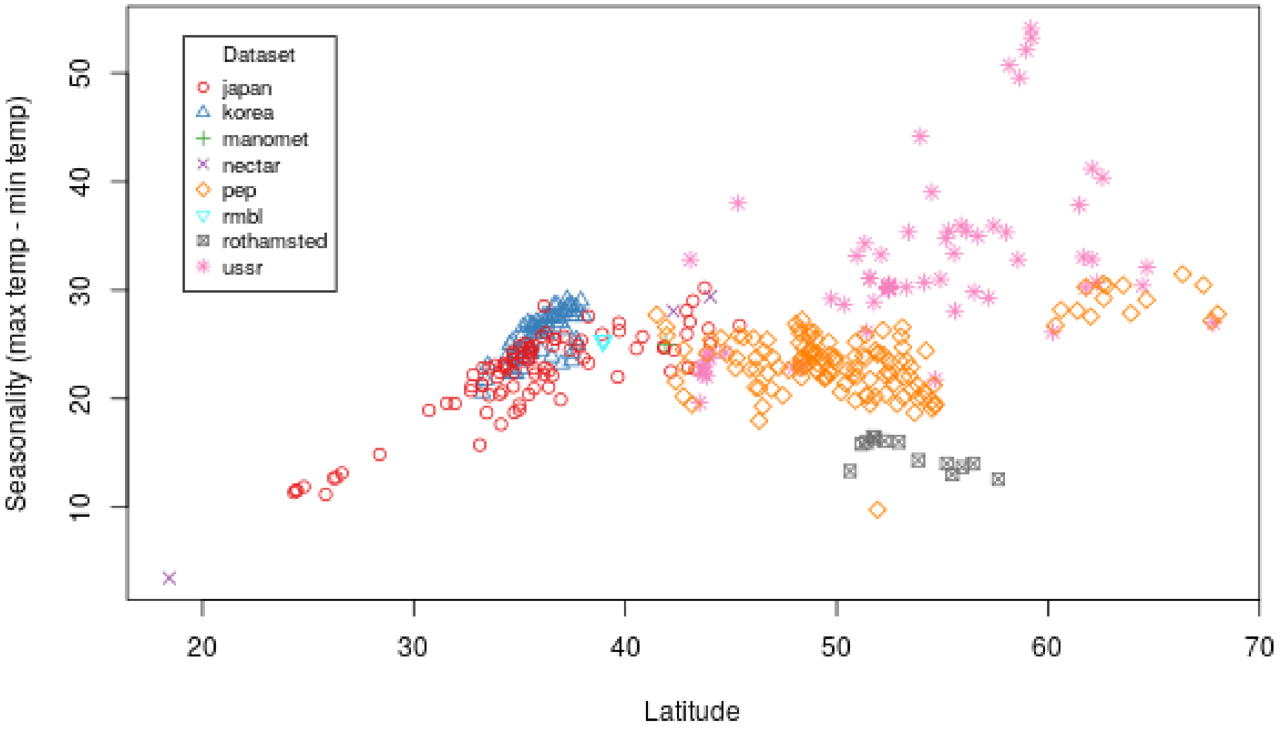

**Figure S6.2**. No one dataset was responsible for the positive relationship of seasonality and mean sensitivity. Each point represents a time-series, colored by the dataset from which it was taken. Lines represent a linear model of mean sensitivity predicted by seasonality with each dataset sequentially withheld. The color of the line corresponds to the dataset that was withheld from the model. Excluding the Rothamsted dataset results in the smallest slope (gray line), and excluding the Chronicles of Nature Calendar dataset (ussr) results in the greatest slope (pink line).

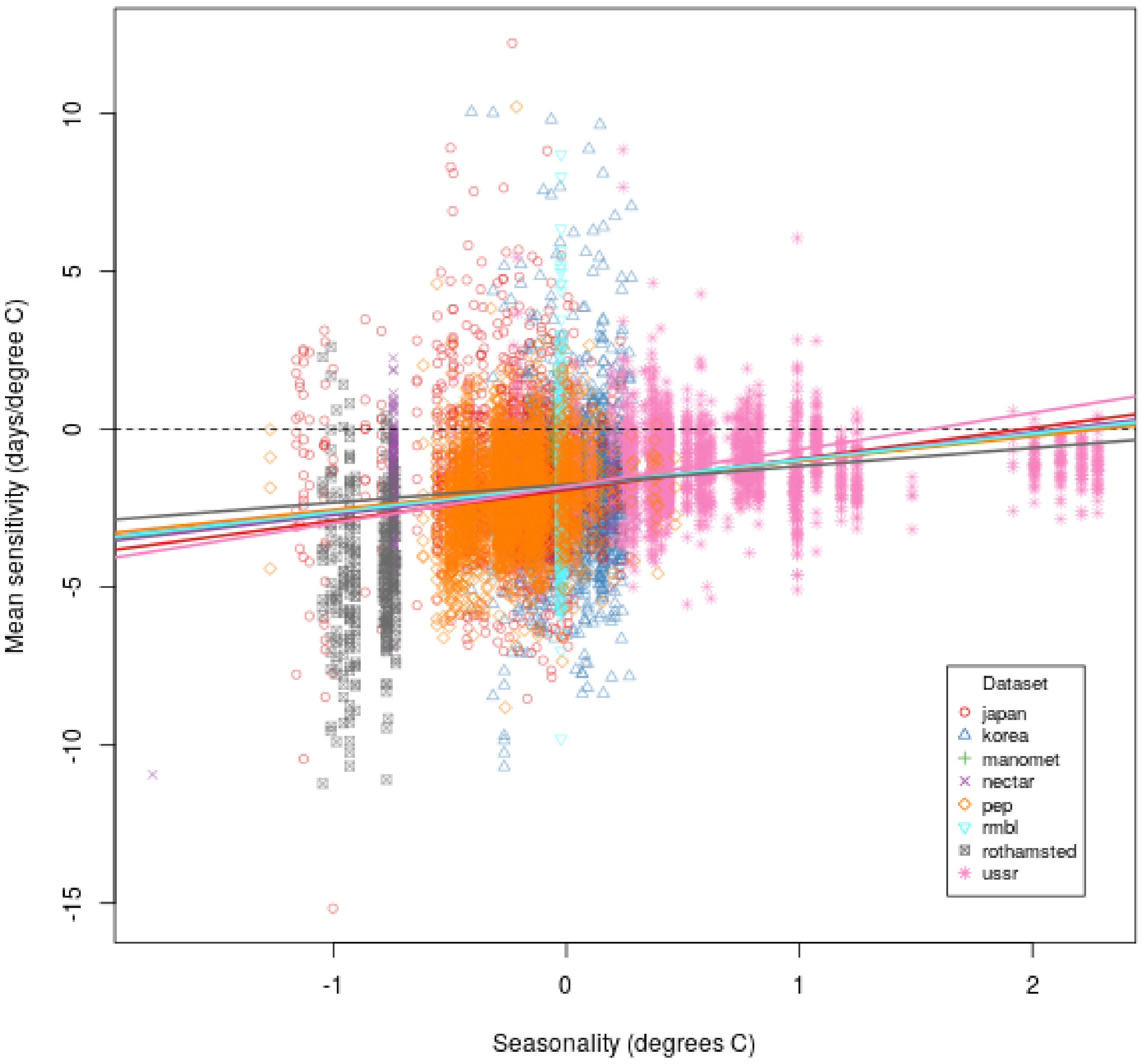

**Figure S6.3**. Changes in population size can artificially shift phenology mean and variance estimates when first-occurrence data is used. As population size increases, the observed first-occurrence advances and variance decreases. For each population size level (n=10, n=1000, n=100000), n draws were made from normal distributions with mean 100 and standard deviation 10, the minimum observation was recorded (points shown) and the procedure was performed 1000 times.

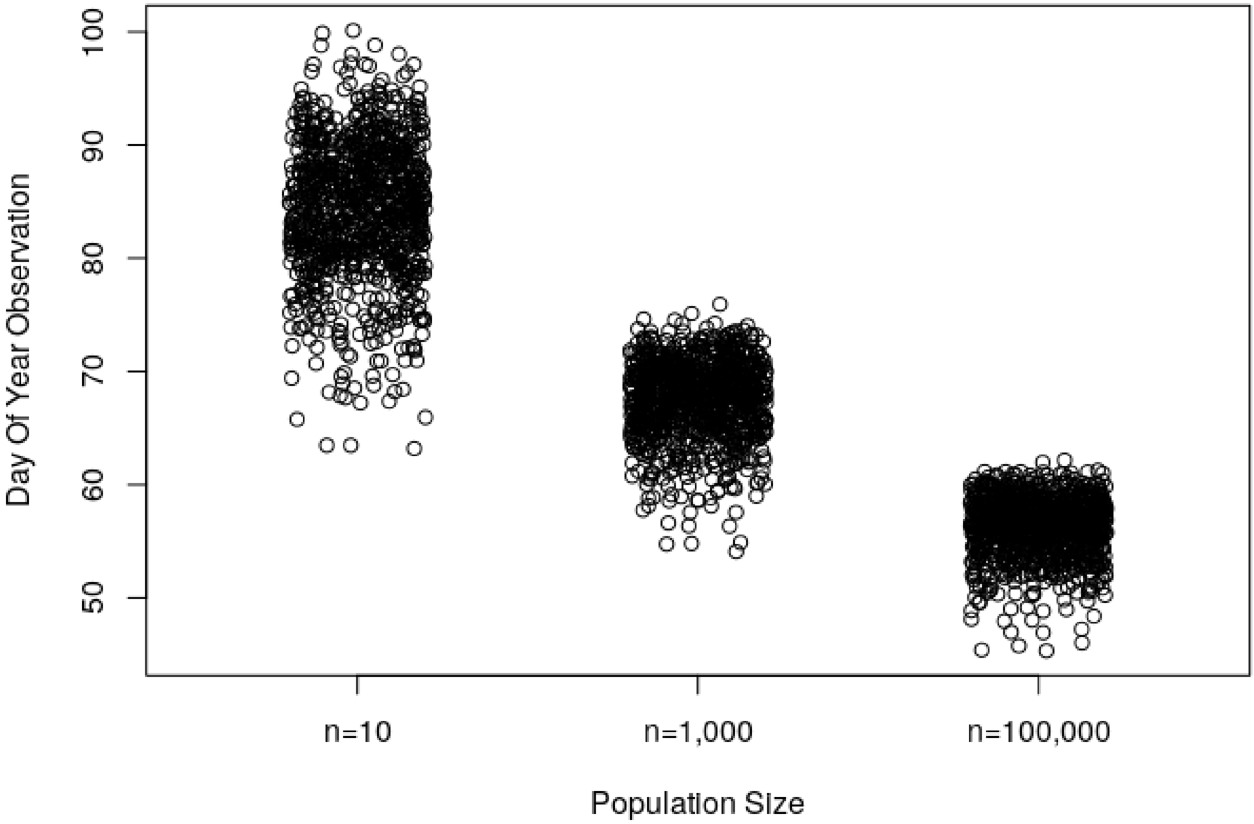

**Figure S6.4**. Seasonal environmental filters may change the skewness of phenological distributions and therefore the inter-annual variance of first-observation dates. In this hypothetical example, there is an environmental filter (red line) in the early season that interacts with the underlying phenological distribution (light blue solid line) to produce a filtered, realized distribution (light blue dashed line). The influence of the environmental filter only comes out when the phenological distribution shifts into its time window; historic, unadvanced phenology (dark blue line) is effectively not influenced by the environmental filter.

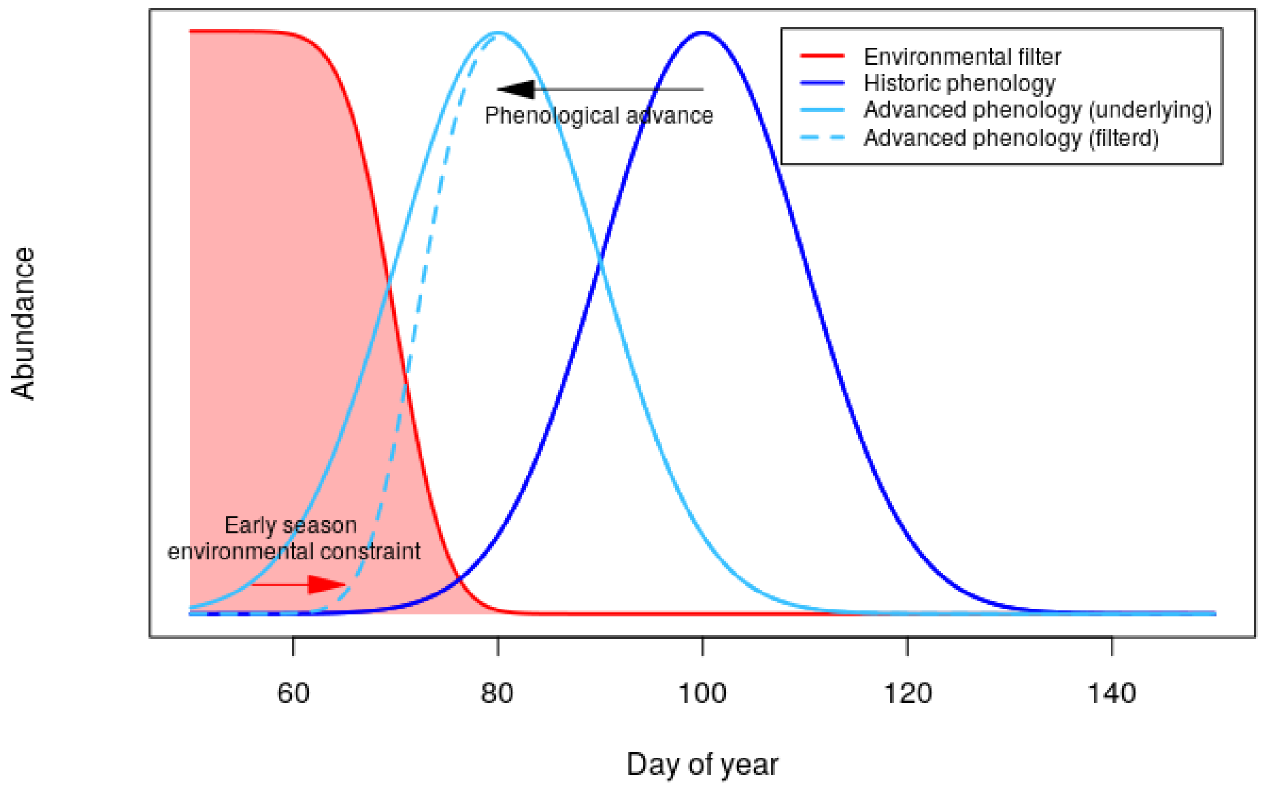

**Figure S6.5**. Spring temperatures have increased markedly at the site locations of this study (left panel), but inter-annual variance has decreased slightly on average (right panel).

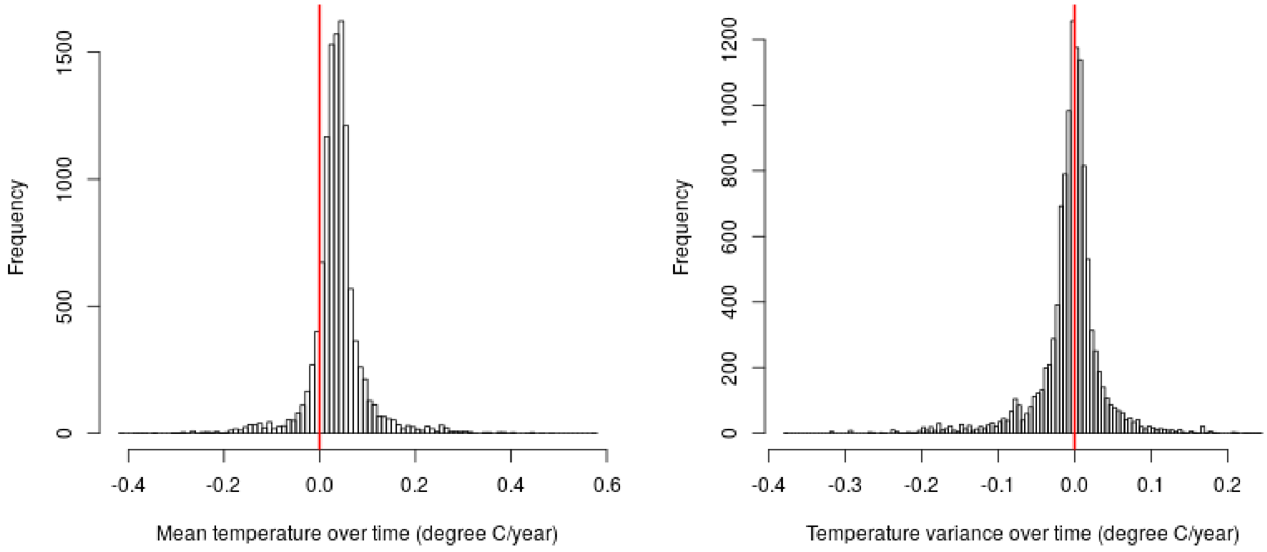

**Supplement 7. Data summary**

**Table S7.1**. Data summary table

| dataset | n time-series | sites | species | n obs. | mean time-series length | mean lat. | mean lon. | mean annual max. temp. | mean seaslty | data range | pheno. groups |
|---|---|---|---|---|---|---|---|---|---|---|---|
| PEP | 3,188 | 134 | 38 | 129,278 | 41 | 50.3 | 12.6 | 13.0 | 22.9 | 1958-2018 | leaves, flowers |
| CNC | 2,278 | 76 | 586 | 69,817 | 31 | 55.5 | 57.8 | 6.8 | 33.7 | 1958-2017 | all |
| Japan | 1,672 | 97 | 39 | 73,760 | 44 | 35.8 | 136.4 | 18.5 | 22.9 | 1958-2011 | all |
| Korea | 1,391 | 68 | 15 | 44,150 | 32 | 36.1 | 127.8 | 17.5 | 26.0 | 1958-2006 | all |
| RMBL | 521 | 30 | 91 | 11,706 | 22 | 39.0 | -107.0 | 8.9 | 25.3 | 1973-2017 | flowers |
| Rothamsted | 337 | 14 | 49 | 11,697 | 35 | 53.3 | -1.5 | 13.2 | 15.0 | 1976-2016 | insects |
| NECTAR | 283 | 4 | 257 | 8,441 | 30 | 49.7 | -14.3 | 13.8 | 18.8 | 1958-2009 | leaves, flowers |
| Manomet | 53 | 1 | 53 | 1,860 | 35 | 41.9 | -70.5 | 14.7 | 25.0 | 1970-2018 | birds |

**Supplement 7. Model coefficient tables**

**Table S7.1.** Main model coefficients. Estimates are standardized effect sizes.

| Metric | Coefficient | Estimate | Std. Error | df | t-value | p-value |
|---|---|---|---|---|---|---|
| μ shift | leaf | -0.163 | 0.011 | 1,034.262 | -14.957 | < 0.001 |
| | flower | -0.166 | 0.010 | 685.227 | -17.164 | < 0.001 |
| | insect | -0.137 | 0.021 | 652.433 | -6.463 | < 0.001 |
| | bird | -0.050 | 0.017 | 1,021.554 | -2.935 | 0.003 |
| | seasonality | 0.153 | 0.019 | 328.303 | 7.919 | < 0.001 |
| | mean_temp | 0.167 | 0.019 | 439.729 | 8.984 | < 0.001 |
| | pheno_position | 0.025 | 0.009 | 1,554.441 | 2.714 | 0.007 |
| μ sensitivity | leaf | -1.639 | 0.055 | 1,188.093 | -29.687 | < 0.001 |
| | flower | -1.823 | 0.049 | 808.816 | -36.916 | < 0.001 |
| | insect | -2.499 | 0.114 | 710.488 | -21.929 | < 0.001 |
| | bird | -1.189 | 0.086 | 917.201 | -13.795 | < 0.001 |
| | seasonality | 1.031 | 0.090 | 401.207 | 11.396 | < 0.001 |
| | mean_temp | 0.611 | 0.091 | 612.461 | 6.741 | < 0.001 |
| | pheno_position | 0.704 | 0.051 | 2,132.067 | 13.845 | < 0.001 |
| σ shift | leaf | -0.028 | 0.005 | 522.782 | -5.363 | < 0.001 |
| | flower | -0.026 | 0.004 | 321.561 | -6.061 | < 0.001 |
| | insect | -0.008 | 0.009 | 269.744 | -0.896 | 0.371 |
| | bird | -0.018 | 0.008 | 457.714 | -2.207 | 0.028 |
| | seasonality | -0.013 | 0.009 | 234.330 | -1.398 | 0.164 |
| | mean_temp | 0.007 | 0.009 | 287.167 | 0.786 | 0.432 |
| | pheno_position | -0.005 | 0.005 | 349.423 | -1.118 | 0.264 |
| σ sensitivity | leaf | -0.247 | 0.027 | 476.377 | -9.219 | < 0.001 |
| | flower | -0.257 | 0.022 | 307.001 | -11.939 | < 0.001 |
| | insect | -0.217 | 0.049 | 329.376 | -4.377 | < 0.001 |
| | bird | -0.155 | 0.041 | 484.145 | -3.809 | < 0.001 |
| | seasonality | 0.165 | 0.046 | 340.480 | 3.576 | < 0.001 |
| | mean_temp | 0.118 | 0.045 | 431.180 | 2.612 | 0.009 |
| | pheno_position | 0.125 | 0.026 | 330.853 | 4.735 | < 0.001 |

**Table S7.2**. Plant traits model coefficients. Estimates are standardized effect sizes.

| Metric | Coefficient | Estimate | Std. Error | df | t-value | p-value |
|---|---|---|---|---|---|---|
| μ shift | (Intercept) | -0.143 | 0.109 | 1,403.583 | -1.310 | 0.19 |
| | grass | -0.170 | 0.116 | 934.409 | -1.472 | 0.141 |
| | herb | -0.123 | 0.109 | 1,333.079 | -1.128 | 0.26 |
| | shrub | -0.060 | 0.113 | 1,047.887 | -0.532 | 0.595 |
| | tree | -0.041 | 0.116 | 1,005.904 | -0.358 | 0.721 |
| | height | -0.114 | 0.052 | 168.314 | -2.181 | 0.031 |
| | seed_mass | -0.037 | 0.028 | 58.147 | -1.301 | 0.198 |
| | SLA | -0.053 | 0.028 | 157.497 | -1.898 | 0.059 |
| μ sensitivity | (Intercept) | -0.973 | 0.591 | 713.670 | -1.646 | 0.1 |
| | grass | -0.275 | 0.648 | 529.494 | -0.424 | 0.671 |
| | herb | -0.871 | 0.589 | 663.596 | -1.479 | 0.14 |
| | shrub | -0.995 | 0.625 | 535.338 | -1.591 | 0.112 |
| | tree | -1.087 | 0.648 | 527.495 | -1.676 | 0.094 |
| | height | 0.016 | 0.332 | 243.254 | 0.047 | 0.963 |
| | seed_mass | -0.173 | 0.199 | 144.745 | -0.870 | 0.386 |
| | SLA | 0.044 | 0.180 | 250.357 | 0.242 | 0.809 |
| σ shift | (Intercept) | -0.010 | 0.073 | 1,657.844 | -0.141 | 0.888 |
| | grass | 0.001 | 0.077 | 1,114.938 | 0.007 | 0.995 |
| | herb | -0.006 | 0.073 | 1,559.870 | -0.084 | 0.933 |
| | shrub | -0.029 | 0.075 | 1,269.201 | -0.381 | 0.703 |
| | tree | -0.015 | 0.077 | 1,225.328 | -0.188 | 0.851 |
| | height | 0.018 | 0.033 | 154.762 | 0.551 | 0.583 |
| | seed_mass | 0.017 | 0.017 | 52.064 | 0.966 | 0.339 |
| | SLA | -0.002 | 0.018 | 144.215 | -0.094 | 0.926 |
| σ sensitivity | (Intercept) | 0.234 | 0.390 | 1,560.609 | 0.599 | 0.549 |
| | grass | -0.773 | 0.405 | 839.014 | -1.907 | 0.057 |
| | herb | -0.572 | 0.388 | 1,534.208 | -1.472 | 0.141 |
| | shrub | -0.407 | 0.397 | 1,149.309 | -1.026 | 0.305 |
| | tree | -0.484 | 0.407 | 962.045 | -1.190 | 0.234 |
| | height | 0.016 | 0.155 | 43.579 | 0.102 | 0.919 |
| | seed_mass | -0.127 | 0.072 | 16.715 | -1.769 | 0.095 |
| | SLA | 0.136 | 0.082 | 42.604 | 1.657 | 0.105 |

**Table S7.3**. Plant phenophase model coefficients. Estimates are standardized effect sizes.

| Metric | Coefficient | Estimate | Std. Error | df | t-value | p-value |
|---|---|---|---|---|---|---|
| μ shift | (Intercept) | -0.179 | 0.024 | 55.643 | -7.513 | < 0.001 |
| | tree | -0.036 | 0.028 | 41.145 | -1.295 | 0.202 |
| | first_leaf | 0.015 | 0.025 | 2,380.520 | 0.598 | 0.55 |
| | tree:first_leaf | 0.008 | 0.027 | 2,346.263 | 0.306 | 0.76 |
| μ sensitivity | (Intercept) | -1.905 | 0.159 | 74.275 | -11.986 | < 0.001 |
| | tree | -0.148 | 0.189 | 63.732 | -0.781 | 0.438 |
| | first_leaf | 0.001 | 0.129 | 2,477.823 | 0.009 | 0.993 |
| | tree:first_leaf | 0.093 | 0.140 | 2,478.481 | 0.661 | 0.508 |
| σ shift | (Intercept) | -0.041 | 0.014 | 41.428 | -2.895 | 0.006 |
| | tree | 0.036 | 0.017 | 33.334 | 2.153 | 0.039 |
| | first_leaf | -0.005 | 0.017 | 2,336.318 | -0.293 | 0.77 |
| | tree:first_leaf | -0.018 | 0.018 | 2,280.007 | -0.967 | 0.334 |
| σ sensitivity | (Intercept) | -0.234 | 0.053 | 51.507 | -4.406 | < 0.001 |
| | tree | -0.072 | 0.060 | 36.484 | -1.215 | 0.232 |
| | first_leaf | 0.052 | 0.084 | 1,756.016 | 0.620 | 0.535 |
| | tree:first_leaf | -0.051 | 0.091 | 1,626.435 | -0.558 | 0.577 |

**Table S7.4**. Bird traits model coefficients. Estimates are standardized effect sizes.

| Metric | Coefficient | Estimate | Std. Error | df | t-value | p-value |
|---|---|---|---|---|---|---|
| μ shift | (Intercept) | 0.026 | 0.024 | 107.894 | 1.073 | 0.286 |
| | Herbivore | 0.053 | 0.033 | 99.980 | 1.577 | 0.118 |
| | Omnivore | 0.010 | 0.031 | 109.233 | 0.320 | 0.75 |
| | log(mass) | -0.047 | 0.025 | 143.837 | -1.880 | 0.062 |
| μ sensitivity | (Intercept) | -0.601 | 0.112 | 81.411 | -5.347 | < 0.001 |
| | Herbivore | -0.208 | 0.172 | 149.318 | -1.211 | 0.228 |
| | Omnivore | -0.193 | 0.157 | 148.513 | -1.230 | 0.221 |
| | log(mass) | -0.016 | 0.120 | 164.004 | -0.132 | 0.895 |
| σ shift | (Intercept) | -0.016 | 0.009 | 30.595 | -1.822 | 0.078 |
| | Herbivore | -0.025 | 0.016 | 48.219 | -1.578 | 0.121 |
| | Omnivore | -0.004 | 0.015 | 62.441 | -0.236 | 0.814 |
| | log(mass) | -0.003 | 0.012 | 98.889 | -0.285 | 0.776 |
| σ sensitivity | (Intercept) | -0.138 | 0.040 | 66.452 | -3.422 | 0.001 |
| | Herbivore | 0.012 | 0.085 | 94.523 | 0.138 | 0.891 |
| | Omnivore | 0.103 | 0.078 | 104.130 | 1.315 | 0.192 |
| | log(mass) | 0.079 | 0.058 | 126.917 | 1.366 | 0.174 |
